# Non-canonically regulated heterochronic expression of Hsp70 drives clonal expansion and invasion in *Drosophila* epithelial tumours

**DOI:** 10.1101/2023.02.02.526918

**Authors:** Abhijit Biswas, Gunjan Singh, Vishal Arya, Subhash Chandra Lakhotia, Devanjan Sinha

## Abstract

Oncogenic stress responses are critical modulators of tumour cell adaptation to hostile microenvironments and clonal dynamics. The evolutionarily conserved Hsp70 chaperone promotes tumour cell survival by suppressing apoptosis and facilitating metastatic progression. However, the temporal order and regulation of its expression in neoplastic tumours remains ill-defined. Here, we show a non-canonical regulation of Hsp70 expression during neoplastic transformation in the *Drosophila* wing epithelium. The malignant *lgl⁴ yki^OE^* MARCM clones exhibited a delayed and progressive induction of Hsp70, tightly correlated with clonal expansion and tissue invasion. Loss of Hsp70 function in developing *lgl⁴ yki^OE^* clones suppressed tumour growth in both larval wing discs and allograft assays, underscoring its significant role in tumour fitness and growth. We identify a non-canonical pathway where ROS-activated JNK signalling drives FOXO-dependent induction of Hsp70, bypassing HSF/Hif1α, to promote tumour growth and invasion. Genetic disruption of Nox or JNK abrogated Hsp70 induction and curtailed tumour expansion. Stress adaptation and malignancy of developing *lgl⁴ yki^OE^* epithelial tumour thus require Hsp70 as a redox-responsive, temporally regulated effector of the JNK–FOXO axis.

## Introduction

Tumourigenesis is defined by presence of a multifarious patterns of cellular diversity, driven by clonal selection and proliferation. These clonal populations progressively diverge genetically, transcriptionally, and proteomically, enabling them to achieve varying levels of fitness(Berenblum and Shubik, 1947, Vogelstein and Kinzler, 2015). This acquisition of the neoplastic phenotype is primarily an adaptive response to intrinsic and extrinsic pressures(Janiszewska, 2020). The adaptive features include numerical and structural variations in the chromatin, epigenetic modifications, inflammation, metabolic reprogramming and single nucleotide variants(Chang et al., 2023, Denk and Greten, 2022, Pavlova et al., 2022, Taylor et al., 2018, Widschwendter et al., 2018, Yizhak et al., 2019).

A prevailing view about the multi-step process of tumour development is that, somatic mutations in a cell provide clonal advantages, facilitating its expansion and the subsequent accumulation of additional genetic and epigenetic lesions. Over time, these mutant cells outcompete neighbouring cells, evolving into a heterogeneous mass of invasive populations(Hanahan and Weinberg, 2011, Merlo et al., 2006). Chromosomal abnormalities, causing loss of tumour suppressors, oncogenic fusions or copy number variations in oncogenes, elicit large scale changes in gene expression profiles. While modifications in epigenetic marks and classical cancer mutations influence tumour evolutionary dynamics, many of these changes are tolerable in normal tissues, raising possibilities of additional factors promoting malignancy(Zhang et al., 2024). A recent pan-cancer analysis by ICGC/TCGA whole genomes consortium indicated that around 5% of the cancer cases did not contain any identifiable driver mutations(Kakiuchi and Ogawa, 2021). An emerging perspective thus posits that mutations alone are insufficient for cancer progression, necessitating other molecular prerequisites(Kakiuchi and Ogawa, 2021). One of the drivers of cancer progression is the common over-expression of the chaperone family of stress proteins(Calderwood et al., 2006, Mosser and Morimoto, 2004), which adapts the cancer cells to stresses originating intrinsically or from the tumour microenvironment. However, their precise consequential implications and regulation remain underexplored.

Hsp70 is highly conserved in prokaryotes and eukaryotes, functioning mainly in the protein quality control mechanisms of cells. It is overexpressed in most cancers and is mechanistically implicated in tumour development by enabling cells to withstand stressors such as nutrient deprivation, hypoxia and low pH(Cao et al., 2019, Huesca et al., 1998, Liu et al., 2021).

Emerging evidences highlight its involvement in dysregulating multiple oncogenic pathways, including receptor tyrosine kinases, Ras, and the Akt-mTOR pathway(Murphy, 2013, Park et al., 2001, Rohde et al., 2005, Song et al., 2001, Vostakolaei et al., 2021). Hsp70 also suppresses apoptosis by preventing the activation of pro-apoptotic signals and formation of apoptosome(Park et al., 2002, Park et al., 2001, Ravagnan et al., 2001, Saleh et al., 2000, Stankiewicz et al., 2005, Arya et al., 2007, Srivastava et al., 2019). Given its extensive role in cancer development and progression, Hsp70 has long been considered a promising therapeutic target(Evans et al., 2010, Patury et al., 2009, Powers et al., 2010). However, a clinically approved protocol remains elusive(Goloudina et al., 2012, Sherman and Gabai, 2015), partly due to limited understanding of its expression patterns in developing tumours, and the regulatory factors involved.

Previous work from our lab had shown elevation of constitutively expressed heat shock proteins in tumour backgrounds except for Hsp70 whose expression was regionally restricted(Singh et al., 2022). The present work identifies the non-canonical temporal expression of Hsp70 as a major driver of the tumour’s proliferative phenotype. Using the inducible genetic models of tumourigenesis in *Drosophila* (Rudrapatna et al., 2012), we could trace the expression of Hsp70 right from the stage of tumour initiation through its development. We observed undetectable Hsp70 levels during the early development of tumour clones. However, its expression increased temporally, with isolated Hsp70-positive cells gradually expanding spatially to most of the clonal area which correlated with enhanced clonal size and invasive phenotypes. Depleting or inhibiting Hsp70 significantly slowed tumour growth, reduced expression of the invasive markers and improved host’s survivability. Interestingly, induction of Hsp70 in tumour clones was independent of the canonical transcription factors HSF and Hif1α (Sima). Instead, the cellular redox state appears to be a critical regulator, with JNK-FOXO axis mediating Hsp70 induction. Our findings provide evidence that Hsp70 acts as one of the causal drivers of neoplastic growth and highlights its redox-dependent regulation which offers new avenues for therapeutic intervention targeting this protein.

## Results

### Temporal pattern of Hsp70 expression correlates with growth of tumour clones

Heat Shock Proteins (HSPs) are known to enable cancer cells tolerate various intra-and extra-cellular stresses faced by expanding cancer cell population and are hence overexpressed across the tumour landscape. Meta-analysis of Hsp70 expression across pan-cancer datasets from the TCGA database revealed elevated levels of the protein in multiple tumour types compared to corresponding normal tissues (Fig S1A). This upregulation correlated with higher tumour grades in specific tumour types (Fig S1B and C), suggesting a role for Hsp70 in progression of these tumours. Moreover, patients with low Hsp70 expression exhibited improved overall survival across several carcinoma types (Fig S1D-G). These data and others in the available literature, however, do not provide information about the temporal order of Hsp70 expression during the tumour’s growth cycle.

In our studies, we took advantage of the MARCM genetic model of epithelial tumourigenesis in *Drosophila* for temporally following the growth of a tumour. In agreement with earlier report form our laboratory(Singh et al., 2022), we confirmed that developmentally expressing HSPs like Hsp60, Hsc70 and Hsp83 were ubiquitously elevated in *lgl^4^ yki^OE^* tumour clones (Fig S1H). In contrast, Hsp70 was not detectable during early stages of establishment of clones but appeared at 72 h after clone induction (ACI), with sporadic cells only expressing Hsp70 in some of the developing tumourous clones (Fig 1A).

**Fig 1.**
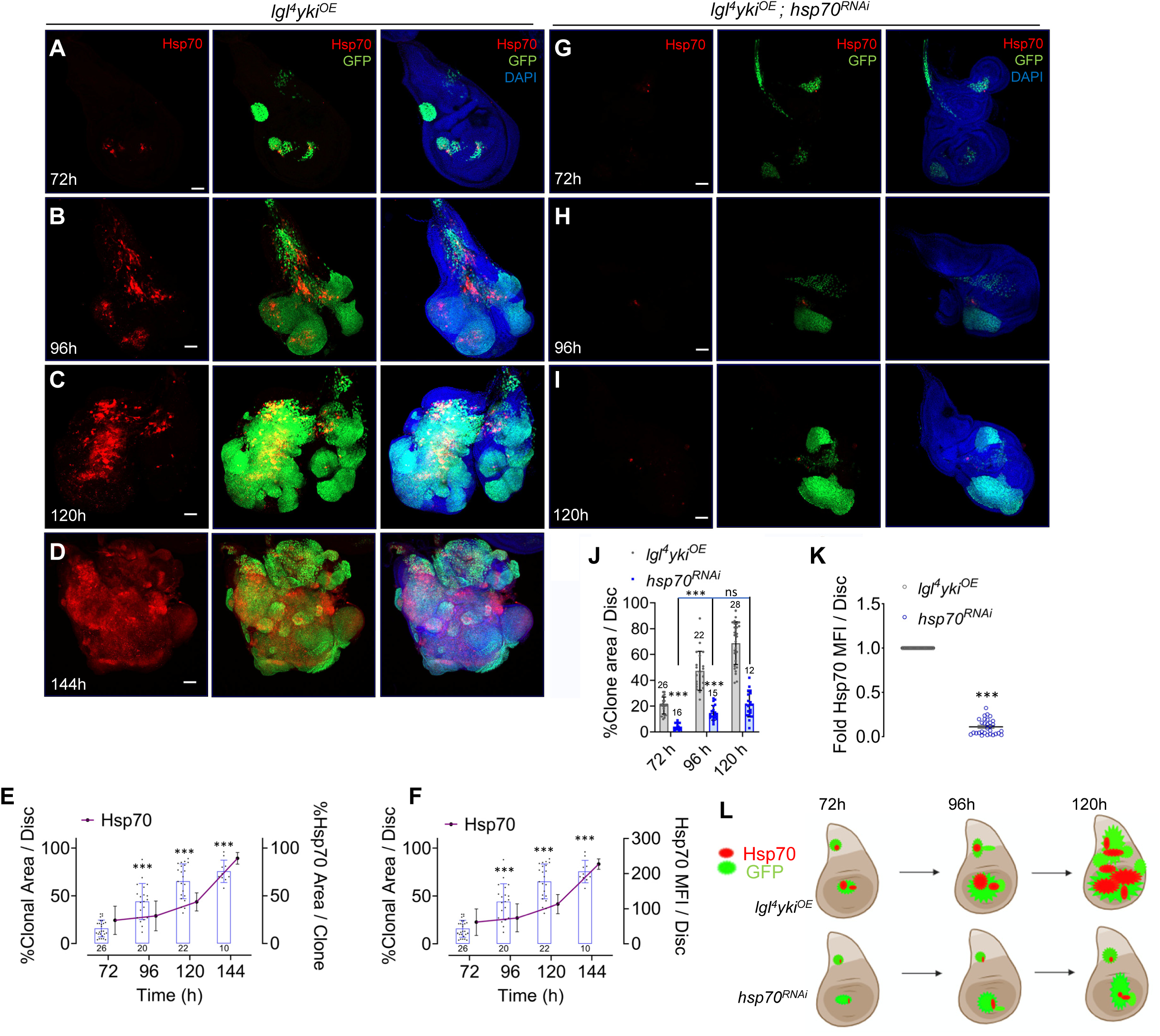
Hsp70 is expressed in a progressive temporal pattern in developing tumours. (**A-D**) Confocal projection images of wing imaginal discs with *lgl^4^ yki^OE^* (green) clones at 72 h, 96h, 120h, 144h ACI, showing stage specific expression and distribution of Hsp70 (red). Scale bar 50 μm. (**E-F**) Dot-bar plots (Y-axis on left) and line graphs (Y-axis on right) showing increased Hsp70 distribution (E) and intensity (F) within clonal area over time. (n = as indicated at the base of each bar). MFI – Mean Fluorescence Intensity. (**G-I**) Effect of *hsp70* downregulation on clonal growth of *lgl^4^ yki^OE^* (green) discs at 72 h, 96h, 120h ACI. Scale bar 50 μm. (**J**) Dot-bar plot showing reduced clonal area in *lgl^4^ yki^OE^*; *hsp70^RNAi^*discs (blue) compared to *lgl^4^ yki^OE^* (gray) across time-points (n = as indicated on top of each bar), corresponding to panels A-G. (**K**) Bar graph showing relative reduction in Hsp70 levels in *lgl^4^ yki^OE^*; *hsp70^RNAi^* wing discs (n = 24) compared to in *lgl^4^ yki^OE^* controls (n = 30) at 96 h ACI. *p < 0.05, **p < 0.01, ***p < 0.001; ns = not significant (p > 0.05); on unpaired Student’s t-test. (L) Schematic summarizing clonal growth and Hsp70 expression in *lgl^4^ yki^OE^* and *lgl^4^ yki^OE^*; *hsp70 RNAi* background.

A progressive increase of Hsp70 expressing cells in *lgl^4^ yki^OE^* tumour clones was noted between 72 to 144 h after the MARCM clone induction (ACI), in parallel with increase in clone size (Fig. 1A-D). At 72 h ACI, only ∼25 % of tumour clones were positive for Hsp70 expression, with the frequency of Hsp70^+^ clones increasing to 30%, 42% and 90% at 96 h, 120 h and 144 h ACI, respectively (Fig. 1E). Correspondingly, the proportion of the GFP⁺ tumour area occupied by Hsp70⁺ cells increased from 15% at 72 h ACI to 80% at 144 h ACI (Fig. 1A-D and I). This was accompanied by a parallel rise in Hsp70 fluorescence intensity and an overall expansion in tumour size (Fig. 1F).

To examine if the above unusual temporal expression of the stress-induced Hsp70 was unique to *lgl^4^ yki^OE^* tumour clones or a common feature in epithelial tumours resulting from different genetic perturbations, we examined Hsp70 expression in other genetically induced hyperplastic or neoplastic tumours. The *lgl^-^* MARCM clones which do not generally become malignant and get eliminated by cell competition with the surrounding wild type neighbours(Khan et al., 2013, Menendez et al., 2010, Nagata and Igaki, 2018), do not show any expression of Hsp70 (Fig. S1I). Likewise, the non-malignant hyperplastic wing discs expressing *MS1096-Gal4* driven *lgl^RNAi^* did not express Hsp70 (Fig. S1J). On the other hand, all other *Act-Gal4* or *MS1096-Gal4* driven malignant tumours (like *Act>lgl^RNAi^, MS1096-GAL4>yki^Act^*, *MS1096-GAL4>Pvr^Act^*, *MS1096-GAL4>Ras^V12^ lgl^RNAi^* or *MS1096-GAL4>Ras^V12^ Scrib^RNAi^*), examined by us showed temporally enhanced Hsp70 expression (Fig. S1K-T). Unlike the *lgl^-^* MARCM clones or the *MS1096-Gal4>lgl^RNAi^*wing discs, the tumours in all the other genotypes displayed high accumulation of F-actin, typically associated with developing neoplasia(Rao et al., 1990) (Fig. S1L-M and S-T). These results indicate that neoplastic tumours having growth advantage express Hsp70 in an incremental pattern comparable to that in *lgl^4^ yki^OE^* MARCM clones. We henceforth, performed all further analysis with wing discs carrying *lgl^4^ yki^OE^* clones, beginning with 72 h ACI which corresponded to early stage of Hsp70 expression.

### Depletion of Hsp70 compromises proliferative and migratory capacity of *lgl^4^ yki^OE^* tumourous clones

Depletion of Hsp70 in *lgl^4^ yki^OE^* tumour clones through co-expression of *hsp70-RNAi* transgene resulted in overall reduction in sizes of *lgl^4^ yki^OE^* clones (Fig. 1G-I and J-K); indicating that expansion of Hsp70^+^ cells paralleled enhanced rate of tumour growth (Fig. 1L). It is noted that compared to a delayed pupation of *lgl^4^ yki^OE^* clone-carrying larvae, *lgl^4^ yk^iOE^; Hsp70 ^RNAi^* larvae pupated by ∼130 h, close to wild-type larval pupation time. This precluded analysis of *lgl^4^ yk^iOE^; Hsp70 ^RNAi^* clones at 144 hr ACI. The enlarged size of *lgl^4^ yki^OE^* tumour clones was associated with high proliferative activity, as evidenced by an increased number of phospho-Histone H3 (pH3^+^) mitotic nuclei in the clonal areas at 72 h (Fig. 2A and E) with higher pH3^+^ cell numbers as the tumour clones progressed to 96 h (Fig. 2C and F). Depletion of Hsp70 led to a reduction in pH3^+^ nuclei that correlated with the observed decrease in clone size. This reduction was consistently at both 72 h (Fig. 2B and E) and 96 h (Fig. 2D and F) clonal stages. RNAi mediated reduction of Hsp70 was also associated with increased expression of the pro-apoptotic marker Dcp1 in 72 h tumourous clones (Fig. 2G, H and K), and an enhanced developmental progression of the tumour carrying larvae. Notably, Dcp1 expression was further increased in Hsp70-depleted tumourous clones at 96 h (Fig. 2I, J and K), indicating that loss of Hsp70 leads to enhanced apoptosis as tumours progress to later stages. Following tumour induction, most larvae form pseudopupae that die without further growth, while some die at the pharate stages, with only ∼50% successfully emerging as adults. Hsp70 downregulation substantially improved their survival so that ∼70% of the *lgl^4^ yki^OE^* clone carrying larvae successfully emerged as adults (Fig. 2L).

**Fig 2.**
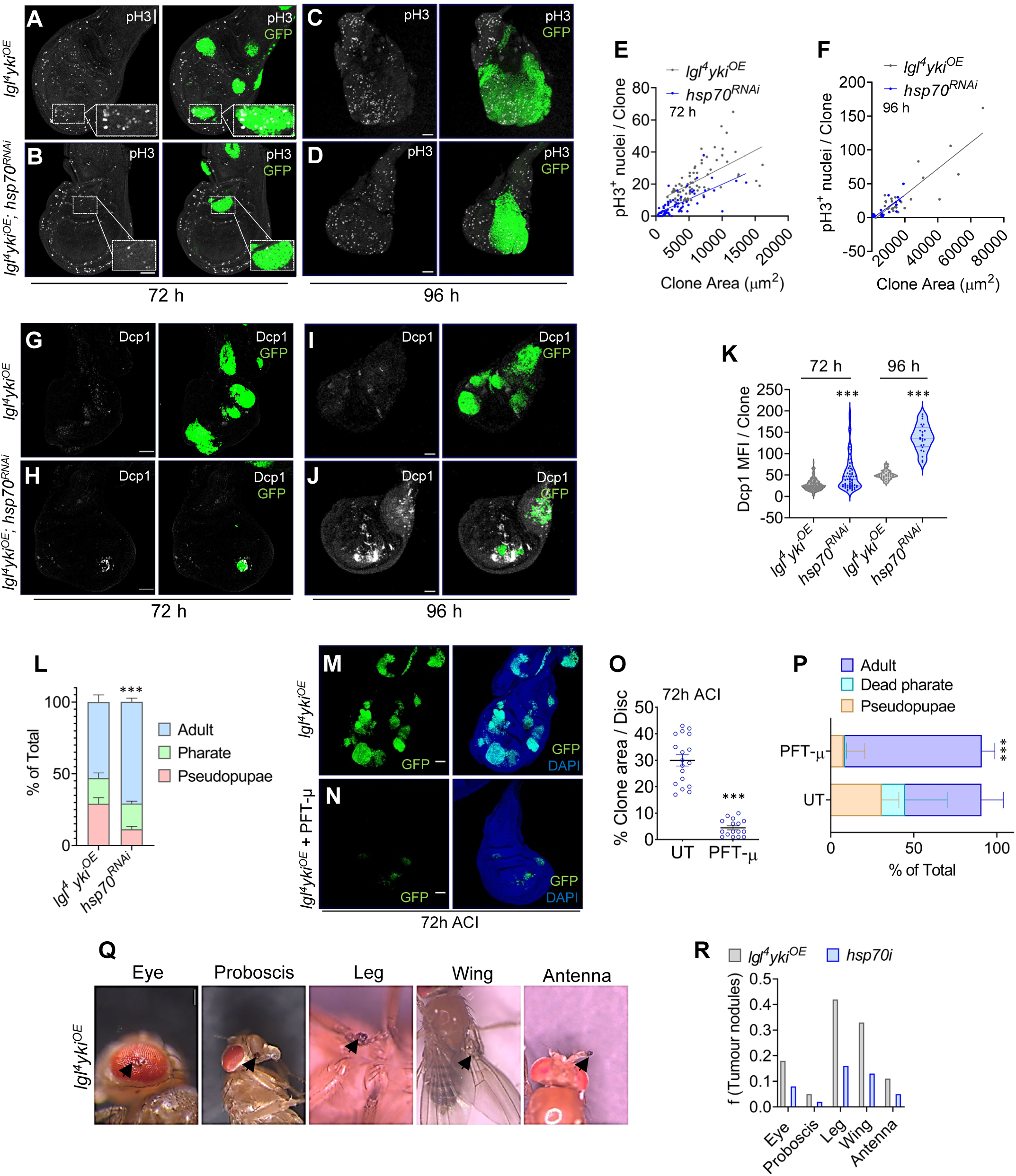
Alterations in Hsp70 levels regress growth and survivability of tumour cells. (**A-F**) Confocal projections of *lgl^4^ yki^OE^* clones (green) at 72 h ACI (A, B) and 96 h ACI (C, D) showing higher numbers of pH3+ mitotic nuclei (grey), compared to *lgl^4^ yki^OE^*; *hsp70^RNAi^*clones. Scale bar: 50 μm. The number of pH3+ foci were quantified and represented as a scatter plot (n = 65 for 72 h and n = 25 for 96 h), p<0.0001 (E, F). (**G-J**) Confocal projections showing Dcp-1 staining (white) within *lgl^4^ yki^OE^* and *lgl^4^ yki^OE^*; *hsp70^RNAi^* clones (green) at 72 h ACI (G, H) and 96 h ACI (I, J), Scale bar: 50 μm. (**K**) The relative Dcp-1 intensity among the samples (G, J) were represented as violin plot (n =146 *lgl^4^ yki^OE^* clones, n = 64 *lgl^4^ yki^OE^*; *hsp70^RNAi^* clones for 72 h and n =45 *lgl^4^ yki^OE^* clones, n = 25 *lgl^4^ yki^OE^*; *hsp70^RNAi^* clones for 96 h), ***p < 0.001, unpaired Student’s t-test. (**L**) Stacked bar graph showing percentages (± S.E.) of pseudopupae, pharate, and adult survivors in *lgl^4^ yki^OE^* and *lgl^4^ yki^OE^*; *hsp70^RNAi^*tumour-bearing backgrounds (n = 200 larvae of each genotype), *p<0.05, **p < 0.01, unpaired Student’s t-test. (**M-O**) Confocal projection images of wing discs showing GFP+ *lgl^4^ yki^OE^*clones in untreated (UT) and 24 h Pifithrin-μ (PFT-μ) treated larval discs at 72 h ACI (Scale bar: 50 μm), further quantified and represented as mean ± s.e.m, n = 18 (UT), n = 16 (PFT-μ), ***p < 0.001 (O). (**P**) Stacked bar graph comparing developmental outcomes (% death as pseudopupae, pharate adults, and adult emergence) between untreated and PFT-μ-treated *lgl^4^ yki^OE^* larvae (n = 50 larvae in each treatment group). (**Q-R**) Frequency of tumour nodules, as shown in the representative image (Q), present in different parts of the surviving adult flies that bore *lgl^4^ yki^OE^*or *lgl^4^ yki^OE^*; *hsp70^RNAi^* tumour clones.

The phenotypic rescue achieved through *hsp70-RNAi* was further confirmed by treating the *lgl^4^ yki^OE^*clone carrying larvae with Hsp70 inhibitor, Pifithrin-μ(Leu et al., 2009), which not only reduced the size of the tumour clones at 72h and 96h ACI (Fig. 2M-O and S2A-C), but also improved the survivability of these larvae as most of them successfully developed to adult stage (Fig. 2P). The observed rescue of *lgl^4^ yki^OE^* clone carrying larvae was not due to reduction in expression of Hsp70 but through reported inhibition of its function by Pifithrin-μ’s binding with the substrate binding domain(Howe et al., 2014) (Fig. S2D-E). Besides the improved survival of *lgl^4^ yki^OE^* clone carrying larvae to adult stage, the frequencies of tumourous nodules in different appendages of adult flies, developing from *lgl^4^ yki^OE^* clone carrying larvae (Fig. 2Q), also decreased following down-regulation of Hsp70 (Fig. 2R).

In addition to diminishing proliferative capacity, Hsp70 depletion also appeared to impair the migratory and invasive properties of tumourous cells. A readout of the aggressive and invasive behaviour is high accumulation of matrix metalloproteinase 1 (MMP1)(Uhlirova and Bohmann, 2006). High levels of MMP1 were detected in 72 h old *lgl^4^ yki^OE^* clones (Fig. 3A, B), which progressively increased with the GFP^+^ clone size at both 96 h and 120 h stages (Fig. 3C-D, E-F). In contrast, MMP1 levels were markedly reduced upon Hsp70 knockdown across examined stages of *lgl^4^ yki^OE^* clones (Fig. 3G-I). Although there was a small increase in size of *lgl^4^ yki^OE^; hsp70^RNAi^* clones at the later stages, this growth was not accompanied by a corresponding increase in MMP1 expression (Fig. 3A’-B’, C’-D’, E’-F’). Pharmacological inhibition of Hsp70 using Pifithrin-μ similarly resulted in a significant reduction in MMP1 levels (Fig. S3A-B and D), along with a comparable reduction is clone size (Fig. S3C). The Hsp70 expressing *lgl^4^ yki^OE^* cells, frequently displayed prominent cytoplasmic edge extensions (Fig. 3J) and a subset of these cells invaded neighbouring tissue (Fig. 3K). Consistently, Hsp70-positive tumour cells showed higher accumulation of disorganized F-actin (Fig. 3L, M), a feature characteristic of migratory and invasive cell populations.

**Fig 3.**
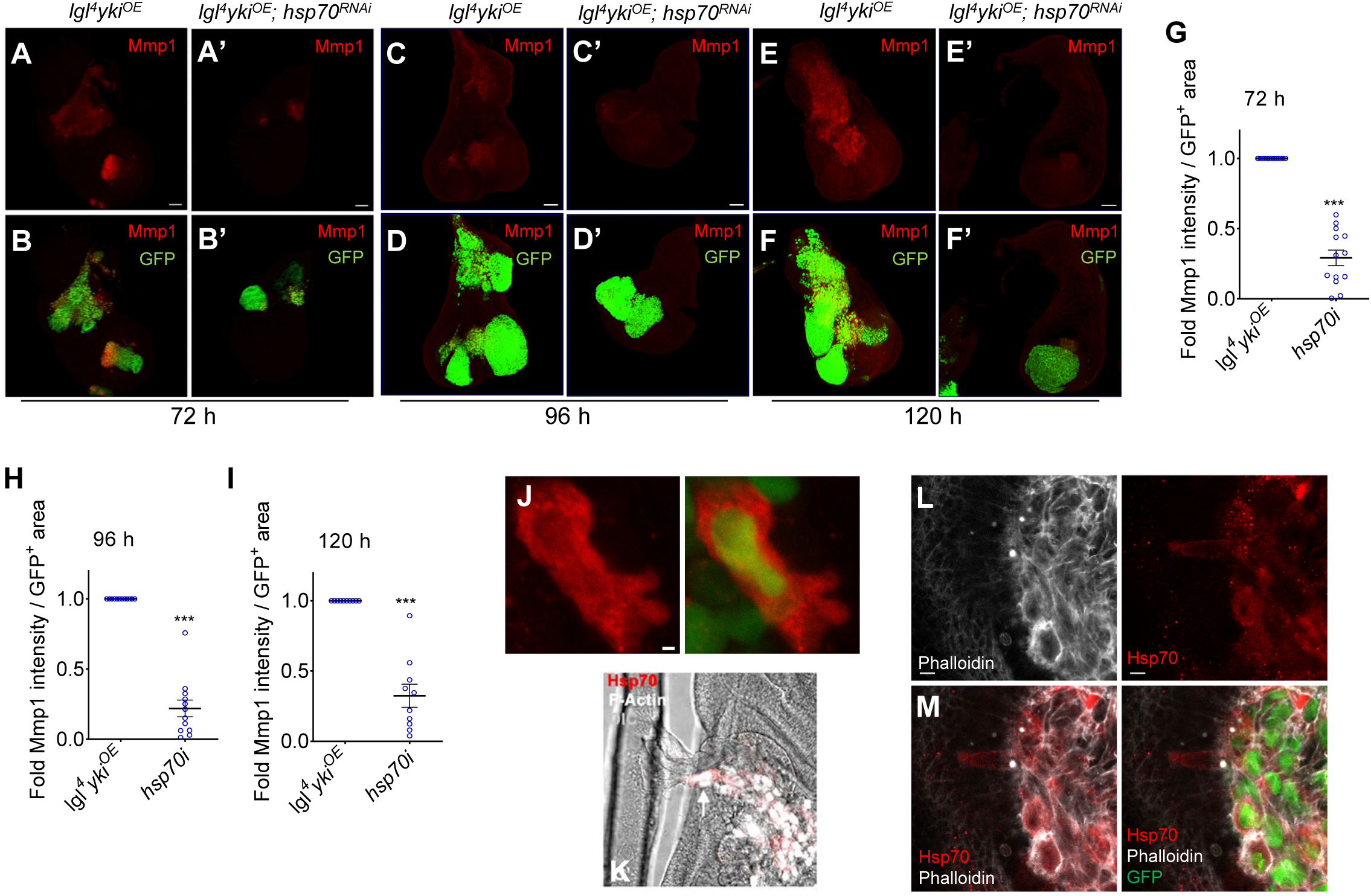
Hsp70 expressing *lgl^4^ yki^OE^* tumour cells show enhanced invasive phenotypes. (**A-I**) Confocal projection images of wing disc bearing *lgl^4^ yki^OE^* and *lgl^4^ yki^OE^*; *hsp70^RNAi^* clones at 72 h ACI (A, B, A’, B’), 96 h ACI (C, D, C’, D’) and 120 h ACI (E, F, E’, F’) showing MMP1 expression within the clonal regions. Scale bar: 50 μm. MMP1 intensity was quantified among size-matched clones and represented as fold change per unit clonal (GFP^+^) area at 72 h (G), 96 h (H) and 120 h (I) after ACI, (n =13 for 72 h, n = 12 for 96 h and n = 10 for 120 h), ***p < 0.001, unpaired Student’s t-test. (**J**) Confocal projection of three optical planes of *lgl^4^ yki^OE^* clones at 96 h ACI, showing altered shape of a GFP^+^ (green) Hsp70^+^ (red) tumour cell, Scale bar: 1 μm. (**K**) Confocal optical section showing Hsp70^+^ clonal cells (red) with high F-actin (white) invading into a neighbouring trachea. (**L-M**) Confocal optical sections depicting accumulation of disorganized F-actin (Phalloidin) in clonal cells with high Hsp70 (M). Scale bar: 2 μm.

### Hsp70 is essential for sustaining tumour growth and invasiveness in allograft tissue

Allografting tumour tissue into a healthy adult *Drosophila* has long served as the gold standard for evaluating tumourigenicity(Miles et al., 2011, Rossi and Gonzalez, 2015, Campbell et al., 2019). To validate the observed correlation between Hsp70 expression and tumour growth, we assessed whether the primary tumours retained similar proliferative and invasive behaviour in an allogeneic environment. Tumour bearing imaginal discs fragments from *lgl**^4^** yki^OE^* and *lgl^4^ yki^OE^*; *hsp70^RNAi^* larvae were dissected at 96 h ACI and transplanted into the abdomens of 2 days old adult wild type (*OregonR^+^*) female flies (Fig. 4A). Post-transplantation, host flies were maintained at 24±1°C and monitored for growth of the transplanted disc fragments. In *lgl^4^ yki^OE^* allografts, the GFP signal from the tumour clones increased progressively from 3 to 14 days assay duration (Fig. 4B and E), indicating sustained tumour expansion. In contrast, *lgl^4^ yki^OE^*; *hsp70^RNAi^* allografts exhibited minimal growth with the growth of GFP foci remaining more or less stunted till 14^th^ day of observation (Fig. 4C and E). The *Nubbin>GFP* wing disc fragments, used as an internal control to monitor the basal growth of the grafted tissue, showed minimal growth (D). The aggressive tumour growth in *lgl^4^ yki^OE^* grafts often resulted in abdominal distension due to fluid accumulation and tumour spread (Fig. 4F and G), whereas *hsp70^RNAi^* grafts did not induce the abdominal inflation of *lgl^4^ yki^OE^*grafts and showed close to normal abdominal morphology similar to flies with *Nubbin>GFP* control disc grafts (Fig. 4F). The proboscis of host fly carrying *lgl^4^ yki^OE^*grafts was swollen and more elongated compared to *lgl^4^ yki^OE^; hsp70^RNAi^* grafted fly (Fig. 4H, I), and this region also showed presence of GFP+ clonal cells (Fig. 4J), indicating possibility of metastasis. Confocal optical section of the host abdomen revealed extensive invasion of *lgl^4^ yki^OE^* tumour cells into nearby tissue and muscles (depicted by dense phalloidin stained F-actin networks) (Fig. 4L-N). Notably, Hsp70-positive clonal cells also showed evidence of distal invasion (Fig. 4M, N). Collectively, these findings reinforce Hsp70 as a critical determinant of tumour clonal growth and invasiveness in the *lgl^4^ yki^OE^* background, and its depletion markedly impairs these malignant properties.

**Fig 4.**
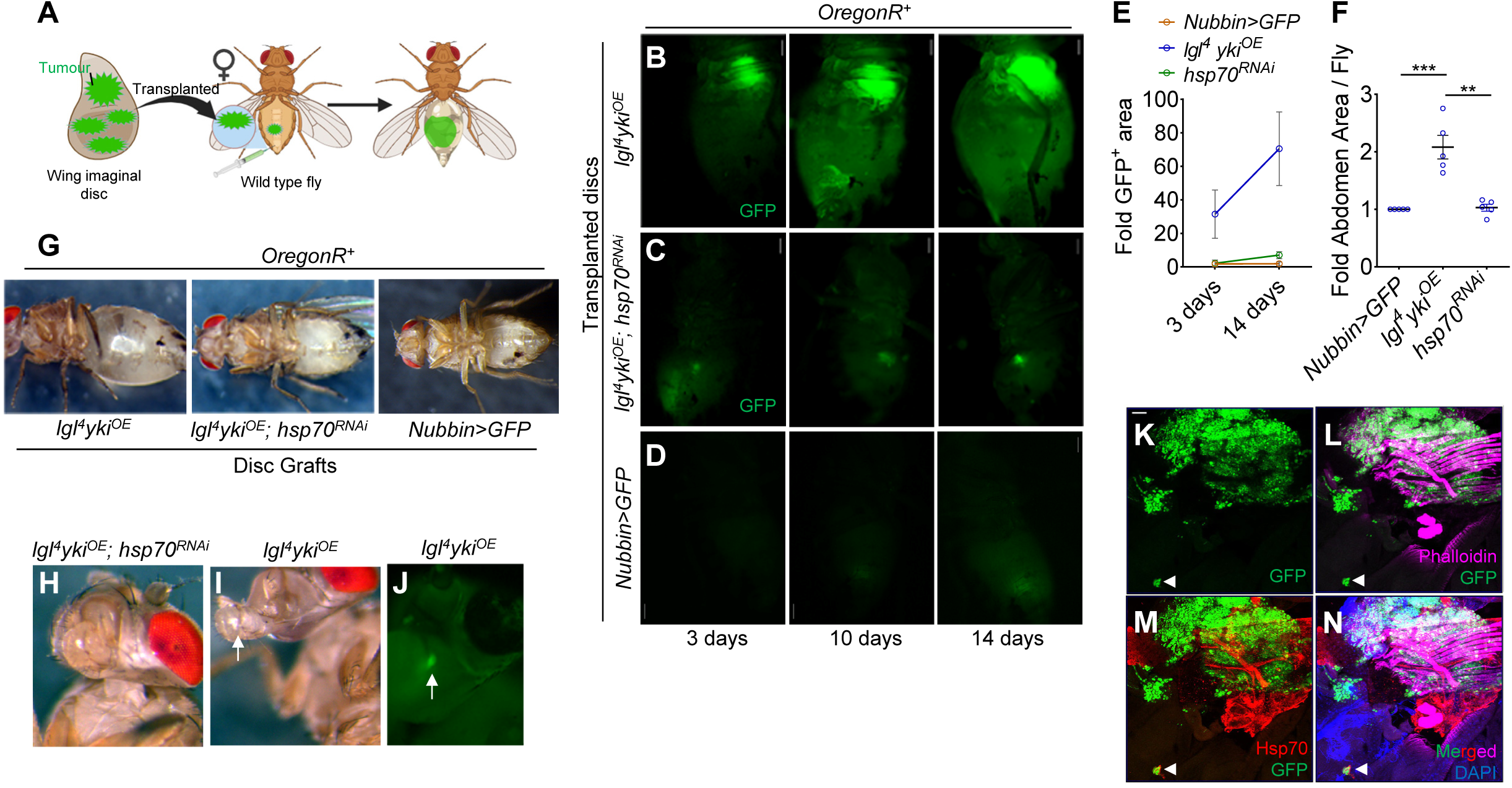
Enhanced invasiveness of *lgl^4^ yki^OE^* tumour allografts in *OregonR^+^* flies is dependent on Hsp70. (**A**) Schematic depicting the wing disc tumour transplantation procedure in *Drosophila*. (**B-D**) Progressive increase in size, indicated by enhancement of GFP fluorescence, of *lgl^4^ yki^OE^* (B) and *lgl^4^ yki^OE^*; *hsp70^RNAi^*(C) disc grafts within the host abdomen over the indicated time periods. *Nubbin>GFP* was used as an internal control (D). (**E**) The relative growth of the disc grafts, as a measure of change in GFP^+^ area, was quantified and represented as fold change over Day 3 *Nubbin>GFP* control. (**F**, **G**) Relative abdominal distension in the host flies (*OregonR^+^*) (G) following grafting of *lgl^4^ yki^OE^, lgl^4^ yki^OE^*; *hsp70^RNAi^* or internal control *Nubbin>GFP* disc fragments. Corresponding abdominal areas shown in panel (G) were quantified and represented as dot plot (F), (n = 5), **p < 0.01, unpaired Student t-test. (**H-J**) Swelling and elongation of proboscis in the host flies carrying *lgl^4^ yki^OE^* allograft (I) and presence of GFP+ clonal cells in the region of proboscis (J), compared to flies carrying *lgl^4^ yki^OE^*; *hsp70^RNAi^*graft (H). (**K-N**) Confocal optical section images at day 14 post-transplantation showing overgrown GFP+ tumour mass (K), local invasion into nearby Phalloidin stained muscle tissue (magenta) (L) and micro-metastases in surrounding tissues marked by GFP^+^ Hsp70^+^ cells (yellow; white arrows) (M, N). Scale bar: 50 μm.

### HSF is not responsible for Hsp70 induction in *lgl^4^ yki^OE^* clones

As a part of evolutionary well-conserved response, heat shock proteins are generally induced in stressed cells by activated transcription factor, HSF(Wu, 1995). Elevated levels of HSF are associated with pro-oncogenic processes in different mammalian tumours and induction of multiple molecular chaperones(Puustinen and Sistonen, 2020). The increased levels of the constitutively expressed heat shock proteins are expectedly HSF independent in developing *lgl^4^ yki^OE^* clones (Singh et al., 2022). In contrast, the stress-inducible Hsp70 appears after establishment of the tumour clones. Therefore, to know if the delayed expression is HSF-dependent, we examined levels of Hsp70 in *lgl^4^ yki^OE^;hsf^RNAi^* background. The HSF expression was relatively similar among *lgl^4^ yki^OE^* clonal and non-clonal cells (Fig. 5A-B and C), and upon *hsf^RNAi^* clone-specific knock-down was achieved (Fig. 5A-B and D-E). Despite effective reduction of HSF in the clones, the magnitude of Hsp70 expression remained unaltered (Fig. 5A-B and F-G) and tumour size showed no significant variation (Fig. 5H). Consequently, co-expression of *hsf^RNAi^* did not rescue the poor survivability associated with *lgl^4^ yki^OE^* tumours (Fig. 5I). These results indicated that Hsp70 expression in these tumours is independent of HSF, implicating possible involvement of alternative non-canonical transcriptional regulators.

**Fig 5.**
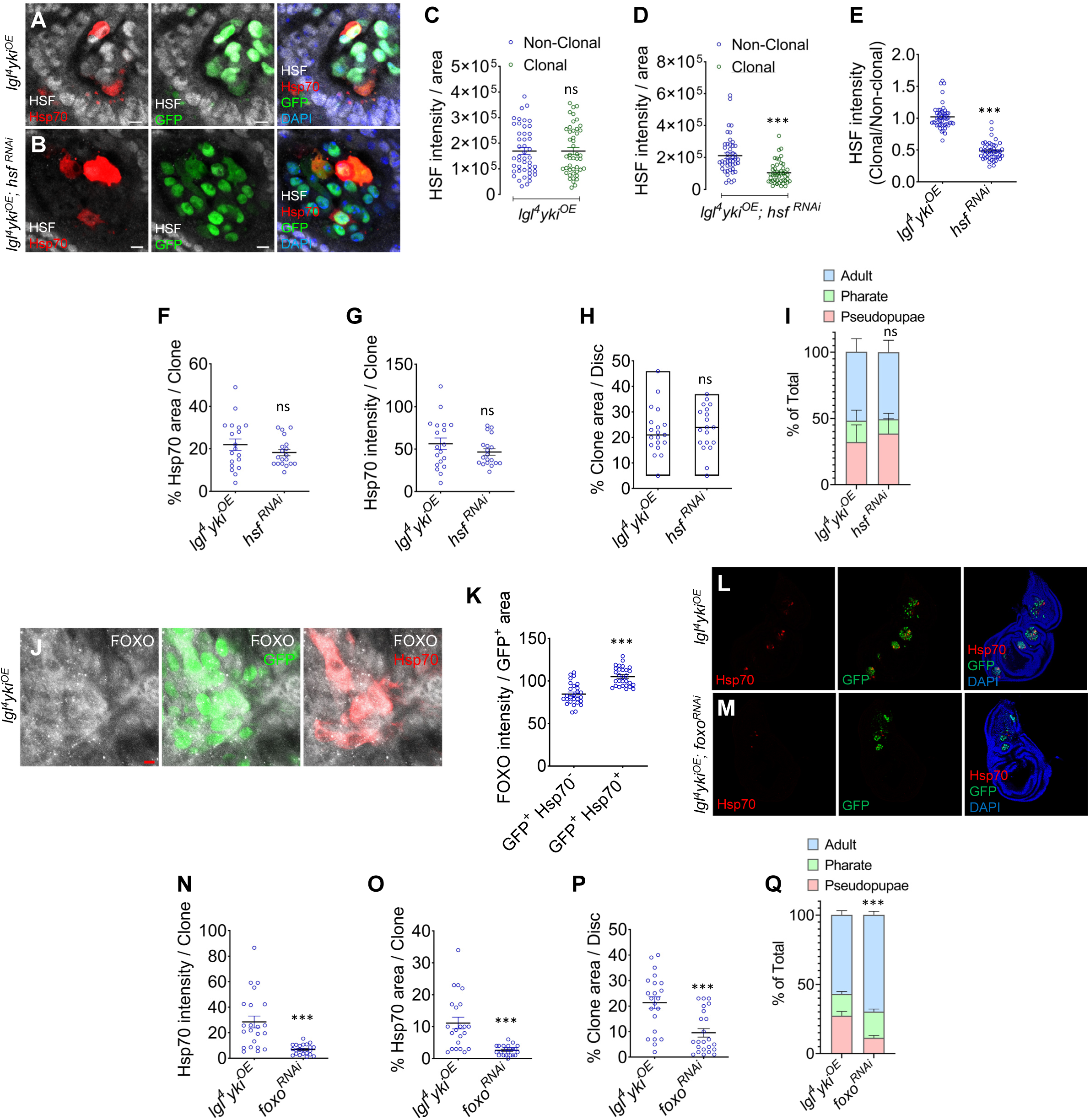
Non-canoncial induction of Hsp70 in *lgl^4^ yki ^OE^* tumours. (**A-B**) Confocal optical sections showing HSF (white) and Hsp70 (red) expression in clonal (GFP^+^, green) and non-clonal (grey) areas of *lgl^4^ yki^OE^* and *lgl^4^ yki^OE^*, *hsf^RNAi^* tumour discs at 72 h ACI. Scale bar: 50 μm. (**C-E**) Quantification of HSF intensity ratio between clonal and non-clonal regions. Ratiometric change in HSF intensity among the clonal and non-clonal area (E), reflected by its unchanged pattern in *lgl^4^ yki^OE^* (C) and significant reduction in its clonal intensity compared to non-clonal regions (D), (n = 46), ***p < 0.001. (**F–H**) Quantification of Hsp70 distribution (F), intensity (G) within clonal regions and the total clonal area (H) between the indicated genotypes, n = 19, ns – not-significant. (**I**) Stacked bar graphs comparing developmental outcomes (pseudopupae, pharate, and adult stages) in *lgl^4^ yki^OE^*, *hsf^RNAi^* compared to *lgl^4^ yki^OE^* (n = 120 larvae), ***p < 0.001. (**J**) Digitally zoomed confocal optical section showing distribution of FOXO and Hsp70 in *lgl^4^ yki^OE^* clones (green) at 72 h ACI. (**K**) FOXO fluorescence intensity in GFP^+^ area (J), which is positive or negative for Hsp70 expression was individually quantified and represented as dot plot, n = 29, ***p<0.001. (**L-M**) Confocal images of *lgl^4^ yki^OE^* and *lgl^4^ yki^OE^; dfoxo^RNAi^* tumour discs showing GFP^+^ clones and Hsp70 expression. Scale bar: 50 μm. (**N-P**) Quantification plots showing Hsp70 intensity, percentage of clonal area covered by Hsp70 and overall clonal size in the indicated genotypes. (n = 22) ***p < 0.001. (**Q**) Stacked bar graphs comparing developmental outcomes (pseudopupae, pharate, and adult stages) in *lgl^4^ yki^OE^; dfoxo^RNAi^* compared to *lgl^4^ yki^OE^* (n = 300 larvae), ***p < 0.001.

### Non-canonical regulation of Hsp70 implicates FOXO as a key upstream driver

Given the prominent role of hypoxia response as one of the major drivers of tumour progression(Mendillo et al., 2012, Scherz-Shouval et al., 2014), we tested the potential involvement of Hif1α (Sima in fruit flies) in induction of Hsp70, since it has a binding site at Hsp70 promoter(Huang et al., 2009). Depletion of Sima in *lgl^4^ yki^OE^* clones by co-expressing *sima-RNAi* transgene, did not cause any significant alteration in Hsp70 expression, its spatial distribution and clonal growth (Fig. S4A-E). Next, we investigated the role of the transcription factor FOXO, proposed to bind the presumptive Forkhead-responsive element (FRE) in Hsp70 promoter(Donovan and Marr, 2016). Notably, Hsp70+ cells within *lgl^4^ yki^OE^* tumour clones displayed high FOXO levels, suggesting a functional correlation (Fig. 5J). The Hsp70^+^, GFP^+^ clonal cells showed stronger expression of FOXO when compared to their non-Hsp70 expressing counterparts (Fig. 5K). Downregulation of FOXO in the *lgl^4^ yki^OE^* tumours resulted in a significant reduction of Hsp70 expression and clonal area (Fig. 5L-P). Correspondingly, *lgl^4^ yki^OE^; foxo^RNAi^* larvae showed better developmental survivability with more than 70% of them emerging into adults (Fig. 5Q). The above observations further supported by reanalysis of publicly available RNAseq data from *Nubbin-Gal4>UAS-yki^S168A^*tumours(Parra and Johnston, 2020) that revealed co-upregulation of *foxo*, *yki*, and *Mmp1*, enrichment of associated signaling pathways, and a concurrent downregulation of *sima* and *hsf* (Fig. S4F-G). These *Nubbin-Gal4>UAS-yki^S168A^* tumours also showed a similar spatial pattern of Hsp70 expression (Fig. S4H). These results indicate that under activated Yki background, expression of Hsp70 is regulated non-canonically by FOXO.

### Redox-mediated JNK–FOXO axis drives Hsp70 expression and tumour expansion

The functional state of FOXO is regulated by upstream kinases-Akt and JNK(Essers et al., 2004, Wang et al., 2005). While Akt suppresses the tumour-suppressive activities of FOXO via phosphorylation and cytoplasmic retention, JNK promotes FOXO activation under stress conditions(Essers et al., 2004). To examine this regulatory dichotomy, we treated *lgl^4^ yki^OE^*tumour-bearing larvae with Capivasertib, a selective Akt inhibitor(Luboff and DeRemer, 2024). Although a marginal increase in clonal size was observed (Fig. 6A-C), Hsp70 expression levels and distribution remained unaffected (Fig. 6A-B, D and E). The phenotypes suggested that FOXO activation and subsequent Hsp70 induction may be largely independent of Akt signaling in this context. To probe the involvement of JNK, we expressed a dominant-negative allele of *basket* (*bsk^DN^*, the *Drosophila* mutant allele of JNK) within *lgl^4^ yki^OE^* clones. Introduction of this allele markedly reduced the number of Hsp70^+^ tumour cells and overall Hsp70 expression (Fig. 6F-G and I-J), which was accompanied by significant clonal growth impairment (Fig. 6H) at 72 h. Since, loss of JNK activity also resulted in a considerable reduction in clone size, we next examined Hsp70 expression at later stages of tumour development, when clones were comparatively larger (Fig. 6K-M), to assess whether reduced Hsp70 levels simply reflected smaller clone size. Notably, even in 96 h tumour clones, Hsp70 expression remained significantly reduced and occupied lesser area per clone (Fig. 6K-L, N-O).

**Fig 6.**
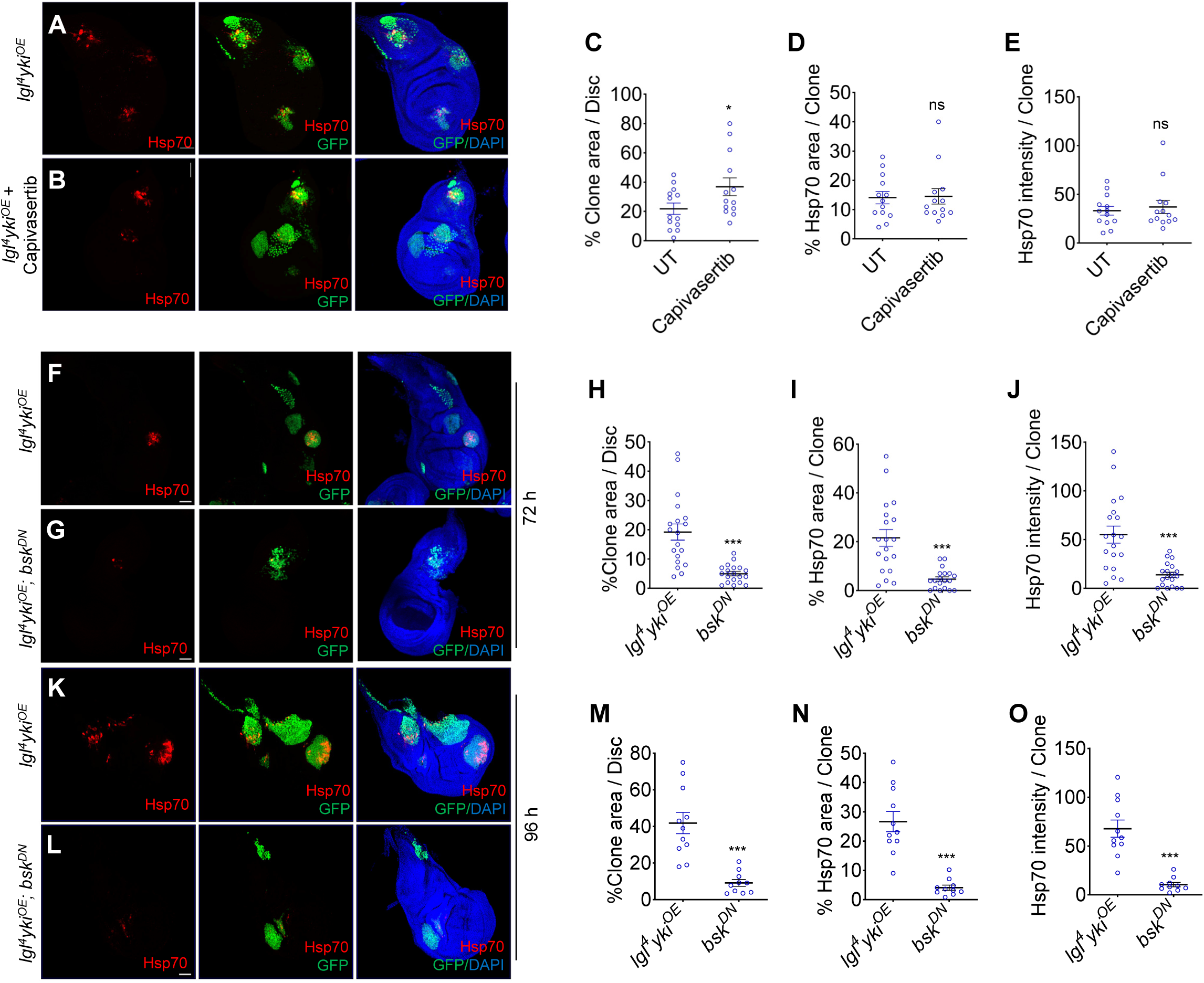
JNK activated FOXO mediates Hsp70 expression. (**A, B**) Confocal projection images showing GFP^+^ clones and Hsp70 expression in untreated (A) and Capivasertib-treated (B) *lgl^4^ yki^OE^* tumourous wing discs at 72 h ACI + 24 h after drug treatment. Scale bar: 50 μm. (**C-E**) Quantification of clonal area in Capivasertib-treated discs (C), Hsp70 distribution (D) and intensity (E) compared to untreated (UT) controls (n = 13). (**F-O**) Confocal projections showing GFP^+^ clonal area and Hsp70 expression in *lgl^4^ yki^OE^* versus *lgl^4^ yki^OE^; UAS-bsk^DN^* tumour discs at 72 h ACI (F, G) and 96 h ACI (K, L). Scale bar: 50 μm. The clonal area, Hsp70 intensity and clonal area covered by Hsp70 was quantified in *lgl^4^ yki^OE^; UAS-bsk^DN^* tumours compared to *lgl^4^ yki^OE^* controls at 72 h ACI (H-J) and 96 h ACI (M-O), (n = 19 for 72 h and n = 10 for 96 h). *p < 0.05, **p < 0.01, ***p < 0.001, ns – non-significant p > 0.05, unpaired Student’s t-test.

As a result, over 80% of larvae harboring *lgl^4^ yki^OE^*; *bsk^DN^* clones successfully developed into adults (Fig. S5A). The observation was further supported by co-localization of elevated pJNK with Hsp70-rich zones within the clonal area (Fig. 7A), underscoring a spatial correlation between JNK activation and Hsp70 induction (Fig. 7B).

**Fig 7.**
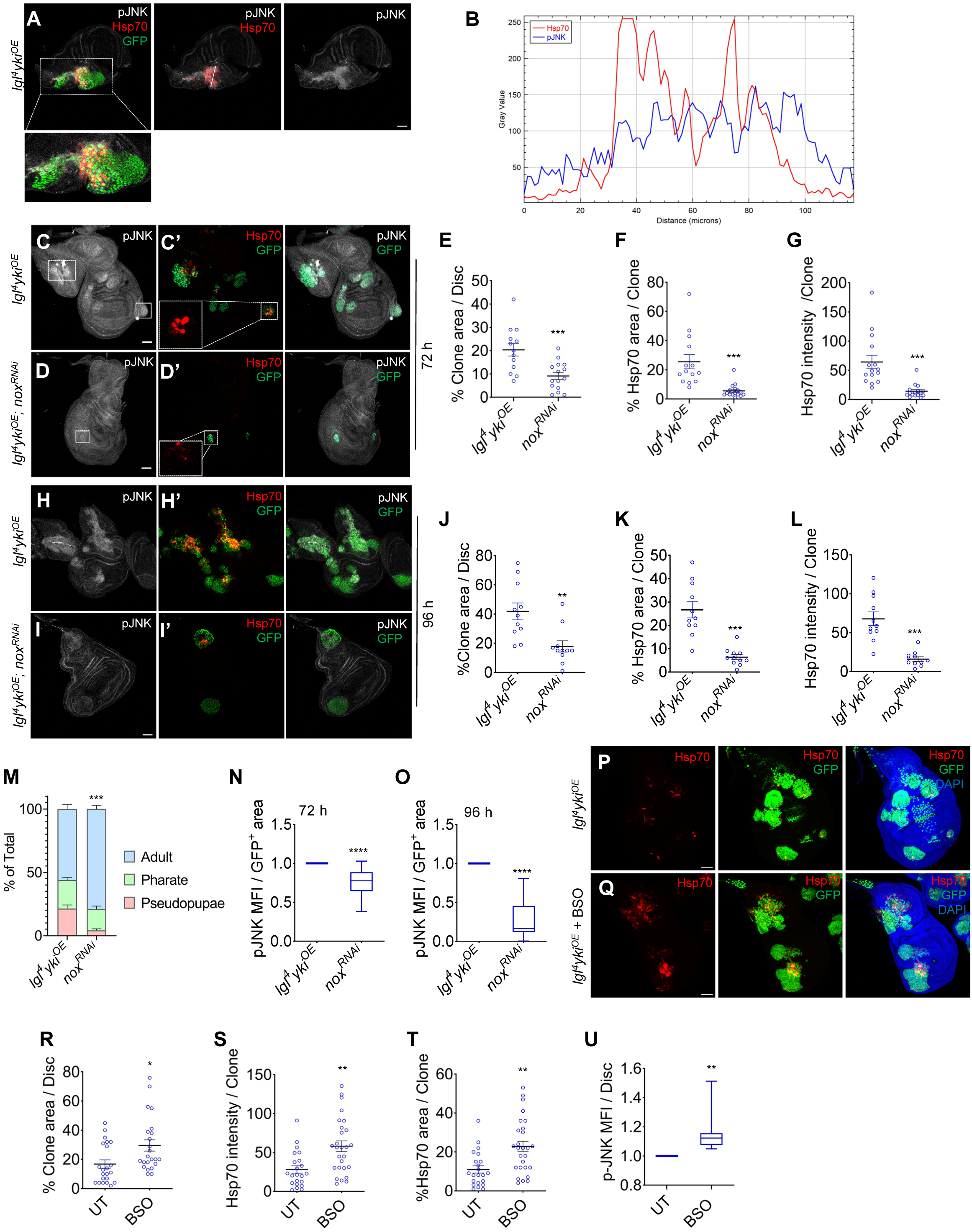
Activation of Hsp70 via FOXO is redox-regulated. (**A, B**) Confocal optical section denoting spatial distribution of Hsp70 expression and activated JNK with the GFP+ clonal area. PlotProfile spatially correlating expression of Hsp70 and JNK (B). (**C-L**) Confocal optical sections showing GFP+ clones and Hsp70 expression in *lgl^4^ yki^OE^* and *lgl^4^ yki^OE^; nox^RNAi^* tumour discs at 72 h ACI (C, D, C’,D’) and 96 h ACI (H, I, H’, I’). Scale bar: 50 μm. The changes in clonal area, Hsp70 coverage of the clonal area and its expression level in *lgl^4^ yki^OE^; nox^RNAi^* tumours (n = 16 for 72 h and n = 11 for 96 h) compared to controls (n = 14 for 72 h and n = 11 for 96 h) was quantified for 72 h (E-G) and 96 h (J-L) ACI discs. (M) Stacked bar graph showing developmental outcomes (pseudopupae, pharate, and adult stages) in *lgl^4^ yki^OE^* vs. *lgl^4^ yki^OE^; nox^RNAi^* tumours (n = 270 larvae of each genotype). (**N, O**) Graph showing changes in pJNK intensity within clonal areas of *lgl^4^ yki^OE^; nox^RNAi^* discs compared to *lgl^4^ yki^OE^* tumours at 72 h ACI (n = 157 clones) and 96 h ACI (n = 11 discs). (**P-Q**) Confocal images of *lgl^4^ yki^OE^* tumour discs showing GFP+ clones and Hsp70 expression in untreated (P) and Buthionine sulfoximine (BSO)-treated (Q) larvae. Scale bar: 50 μm. (**R-T**) Plots showing changes in clonal area (R), Hsp70 expression (S) and its distribution within the clones (T), in BSO treated *lgl^4^ yki^OE^* larvae (n = 27) over untreated (UT) controls (n = 22). (**U**) pJNK intensity in BSO-treated *lgl^4^ yki^OE^* tumours compared to untreated controls (n = 62 clones). *p < 0.05, **p < 0.01, ***p < 0.001, ****p < 0.0001, unpaired Student t-test. Representative image in shown in Fig S6A.

To identify upstream regulators of JNK activation and subsequent FOXO-driven Hsp70 expression and tumour growth, we assessed the role of redox state which is known to have a bearing on tumour growth. NADPH oxidase (Nox), a major source of reactive oxygen species (ROS), is often dysregulated in various tumours, promoting malignant behaviour(Xiong et al., 2025). Therefore, we down-regulated Nox via RNAi in *lgl^4^ yki^OE^* background. The *nox^RNAi^*expressing clones exhibited substantially reduced expression of Hsp70 (Fig. 7C’-D’ and F-G) and impaired clonal growth (Fig. 7E) at 72 h. This reduction in Hsp70 expression was not dependent on the stage of tumour clones. Even at 96 h, *nox^RNAi^* expressing clones continued to display diminished Hsp70 expression, reduced Hsp70-positive clonal coverage, smaller tumour clonal size (Fig. 7H-L). As a result, *lgl^4^ yki^OE^, nox^RNAi^* larvae showed improved survival to adult stage (Fig. 7M). The *lgl^4^ yki^OE^, nox^RNAi^* clones also showed lower levels of pJNK (Fig 7N), with the decrease in pJNK expression becoming more pronounced at later stages of tumour development (Fig. 7O), indicating possible ROS mediated activation of the protein. Notably, under *nox* RNAi conditions, loss of JNK activity was accompanied by a near-complete loss of Hsp70 expression, indicating a requirement for JNK in the regulation of Hsp70 expression (Fig. 7H-I, H’-I’). The Glutathione homeostasis, that acts as a redox buffer, has been observed to be a key regulator of JNK activation(Adler et al., 1999, Wilhelm et al., 1997, Okamura et al., 2015). Treatment with buthionine sulfoximine (BSO), an inhibitor of GSH synthesis, elevated pJNK and Hsp70 levels, which colocalized within tumour clones (Fig. S6A-B, 7U) and corresponded with proportionate expansion of tumour clones (Fig. 7P-T). Conversely, supplementation with N-acetylcysteine (NAC), a precursor of GSH(Lavoie et al., 2008) and a potent antioxidant, led to marked reduction in clonal size and downregulation of Hsp70 expression pattern (Fig. S6C-G). Collectively, these finding demonstrate redox-sensitive activation of Hsp70 through JNK-FOXO axis that result in gradual expansion of tumour clones.

## Discussion

In the competitive tumour landscape, dynamic changes in gene expression profiles determine tumour growth and metastasis. Besides transcription regulators, certain proteins act as drivers of tumour development. Our results suggest that the stress-inducible Hsp70 may play such role in epithelial tumours of *Drosophila.* The MARCM model used by us to generate tumourous clones in developing wing discs, permitted tracing the growth of these clones since their birth(Singh et al., 2022); thus allowing the observation that unlike the constitutively expressed Hsp83/90, Hsc70 and Hsp60, the stress-inducible Hsp70 which is also known to be widely expressed at high levels in various human malignant tumours(Murphy, 2013), appears in tumourous clones only after their establishment, i.e. 48 h to 72 h ACI. The coincidence of expression of Hsp70 in a few cells of the tumour clones during early stages and its subsequent expansion, together with MMP1 accumulation, cytoplasmic protrusions and neighbouring tissue invasion, indicate a causal relationship between Hsp70 and tumour invasiveness. This is also supported by our finding that the neoplastic tumours of different genetic backgrounds show a temporal pattern of Hsp70 expression.

The significantly reduced growth of the *lgl^4^ yki^OE^*clones *in situ* or after allografting in adult flies following either RNAi-or inhibitor-directed down-regulation of Hsp70 activity also supported a role of Hsp70 in driving the developing tumours. Knock-down of Hsp70 also rescues the developmental experienced by larvae carrying *lgl^4^ yki^OE^* clones and improved their survival to adult stage. Our results with *Drosophila* model are also in agreement with an earlier report that the stress inducible Hsp72 expression in malignant mouse mammary tumours is upregulated initially only in a few cancer initiating cells that undergo rapid metastasis(Gong et al., 2015). The stress-inducible Hsp70’s role in cell mobility is also supported by the essential requirement of Hsp70 during the normal developmental migration of border cells in *Drosophila* ovary(Cobreros et al., 2008).

The expression of Hsp70 in tumours is prognostically significant since a positive correlation has been demonstrated between the protein and patient survivability(Ciocca et al., 1993, Sherman and Gabai, 2015, Syrigos et al., 2003). Several studies have proposed a role of Hsp70 in malignant transformation through prevention of apoptosis, modulation of inflammatory and oncogenic pathways (Murphy, 2013, Vostakolaei et al., 2021, Zhao et al., 2023, Sherman and Gabai, 2015). Cancer cells depleted for Hsp70 presented senescent morphological features and cell cycle arrest(Rohde et al., 2005). Hsp70’s pro-tumourigenic function may derive from its interactions with several factors associated with cell growth. Hsp70 plays a positive role in regulating Ras-Erk activation and Hsp70 family members are thought to indirectly promote Erk expression(Cao et al., 2019, Song et al., 2001). Haploinsufficiency or knock-down of Hsp70 reduced KRas levels and caused suppression of lung and pancreatic tumours (Rangel et al., 2021, Shen et al., 2017). Hsp70 members in general are known to facilitate PI3K/Akt/mTOR, promoting cancer progression(Fu et al., 2008, Liu et al., 2021, Liu et al., 2009, Xu et al., 2019). Downregulation of Hsp70 activity from the tumour clones might have lowered the active forms of these factors and made the cells more prone to apoptosis.

Our study provides insights on the regulatory dynamics of Hsp70 in tumour growth using the *Drosophila* tumour models that has an advantage of being genetically stable with relative homogeneity(Gong et al., 2021). Expression of Hsp70 is normally driven by activated HSF following exposure of cells to stress. Particularly in cancer cells, induction of HSF is important in maintaining high levels of molecular chaperones required to counter to proteomic imbalance brought about by genomic alterations and stress(Liao et al., 2015, Kourtis et al., 2018). In addition, HSF is also reported to activate transcriptional programs associated with malignant transformation (Mendillo et al., 2012, Scherz-Shouval et al., 2014). HSF is known to modulate the expression of Hif1α, the master regulator of cellular response to hypoxia, that also induces Hsp70 expression(Chen et al., 2011, Forsythe et al., 1996). Contrary to the known role of HSF as the major regulator of stress-induced Hsp70, our results indicate that expression of Hsp70 in *lgl^4^ yki^OE^* tumours to be independent of both HSF and Hif1α (Sima), since depleting either of the proteins did not affect Hsp70 expression and tumour growth. On other hand, the characteristic expression pattern of Hsp70, that is essential for the tumour fitness, was found to be dependent on FOXO in *lgl^4^ yki^OE^*tumours and is regulated by the cellular redox state. FOXO family of transcription factors are associated with multiple cellular processes such as cell cycle arrest, apoptosis, autophagy, DNA repair, through which it exerts its tumour modulatory effect(Greer and Brunet, 2005, Zhang et al., 2011). Under oxidative stress, JNK phosphorylates FOXO, leading to its activation and nuclear translocation(Essers et al., 2004, Wang et al., 2005). In ischemic mouse models, FOXO physically interacts with Yki (YAP) to regulate antioxidant defence(Shao et al., 2014). In *scrib Ras^V12^*model, lowering of ROS associates with reduction of tumour clones(Perez et al., 2017). Loss of epithelial polarity regulators such *scrib, dlg, lgl* promote tumour through JNK under suppressed Hippo pathway(Enomoto and Igaki, 2011). Reduction of JNK activity, either by its depletion or by changing levels of reduced glutathione, resulted in loss of Hsp70 levels and concomitant reduction in tumour clones. These results were also phenocopied upon downregulating one of the cellular ROS generators, NADH oxidases, or upon exposure to antioxidants. The cell growth driven by oxidative stress-mediated FOXO-dependent activation of Hsp70 supports the dual role for FOXO in regulating both cell proliferative and death pathways. However, we cannot fully exclude the possibility that attainment of a critical clonal size also influences Hsp70 induction, as disruption of the upstream regulators is accompanied by a substantial reduction in clone size.

This study highlights the heterochronicity of Hsp70 expression among developing tumours, emphasizing on its role in enhancing cellular growth and sustenance within the tumour microenvironment. While other molecular chaperones are generally ubiquitously expressed across tumour landscapes(Kourtis et al., 2018, Mosser and Morimoto, 2004), the temporal heterogeneity in Hsp70 expression observed here suggests this chaperone’s unique functional requirements during tumour expansion. Our study raises several compelling questions: what clonal advantages are conferred by Hsp70 that enable certain cells to outcompete their neighbours? While reactive oxygen species (ROS) appear to be the key in inducing Hsp70, what determines its activation in only a subset of tumour cells? Do Hsp70-positive populations exhibit greater immune evasion or possess distinct metabolic traits, and is Hsp70 expression a driver or a consequence of these characteristics? While these questions remain open, our findings offer valuable insights into the expression dynamics of this critical chaperone in tumours. These findings pave the way for exploring innovative strategies for diagnosis and prognosis in mammalian systems, with potential implications for understanding and targeting tumour progression.

## Materials and Methods

### *Drosophila* stocks and genetics

All the fly stocks and crosses were maintained on standard fly food made up of yeast, sugar and agar-maize powder at 24 ± 1°C. The following fly stocks were used to set up crosses to obtain progeny of the desired genotypes:

1. *y 1.* w* *hsFLP tubGAL4 UAS-GFP; tubGAL80 FRT40A/CyO-GFP; +/+;* (Kind gift from Prof. Pradip Sinha, IIT Kanpur).
2. *w*; lgl^4^ FRT40A yki^OE^ /CyO-GFP; +/+* (*lgl* mutant is referred to as *lgl^4^* and *UAS-yki* referred to here as *yki^OE^*); (Kind gift from Prof. Pradip Sinha, IIT Kanpur).
3. *y 1 sc* v 1 sev21*; P{y[+t7.7] v[+t1.8]=TRiP.GLV21028}attP2, (BDSC-35663; referred to as *hsp70^RNAi^*).
4. *y1 v1; UAS-hsf-RNAi/TM3, Sb1*, (BDSC-27070, referred to as *hsf RNAi);*
5. *sc[*] v [1] sev[21]*; P{y[+t7.7]v[+t1.8]=TRiP.HMS00793}attP2 (BDSC-32993, referred to as *dfoxo^RNAi^*).
6. *y[1] sc[*] v[1] sev[21]*; P{y[+t7.7] v[+t1.8]=TRiP.HMS00691}attP2 (BDSC-32902, referred to as *nox^RNAi^*).
7. *y[1] sc[*] v[1] sev[21]*; P{y[+t7.7] v[+t1.8]=TRiP.HMS00833}attP2 (BDSC-33895, referred to as *sima^RNAi^*).
8. *w[*]*; P{w[+mC]=UAS-bsk.K53R}20.1a (BDSC-9311, referred to as *bsk^DN^*)
9. *w[1118]* P{w[+mW.hs]=GawB}Bx[MS1096] (BDSC-8860) referred to as MS1096-Gal4
10. *y[1] w[*];* P{w[+mC]=Act5C-GAL4}25FO1/CyO, y[+] (BDSC-4414) referred to as *Act-Gal4*
11. *w1118; +/+; UAS-RasV12* (BDSC-4847): This transgenic construct is inserted on chromosome 3 which expresses activated Ras protein, under UAS promoter, with a single point mutation in Ras sequence resulting in Glycine at 12 position being replaced by Valine.
12. *w1118; UAS-RasV12/CyO; +/+:* This is a kind gift from Prof. Pradip Sinha (IIT Kanpur). It expresses activated RasV12 protein under UAS promoter; the transgene is inseted on chromosome 2.
13. *y1 sc* v1 sev21*; P{y[+t7.7] v[+t1.8]=TRiP.HMS01490}attP (BDSC-35748): This is referred to here as scrib-RNAi.
14. *y1 sc[* v[1 sev[21*]; P{y[+t7.7] v[+t1.8]=TRiP.HMS01522}attP2 (BDSC-35773): This is referred to here as lgl-RNAi
15. *w*; +/+; P{y[+t7.7]* w[+mC]=UAS-yki.S111A.S168A.S250A.V5}attP2 (BDSC-28817): This transgene is inserted on 3rd chromosome and expresses V5-tagged Yorkie protein with S111A, S168A and S250A amino acid substitutions under UAS control. This stock is referred to here as UAS-yki^Act^.
16. *w[1118]*; P{w[+mC]=UASp-Pvr.lambda}mP1 (BDSC-58428) reffered to as Pvr^act^
17. *OregonR+*: Wild type strain of *Drosophila melanogaster*, used as host in tumour transplantation experiment.
18. w[*]; P{w[nub.PK]=nub-GAL4.K}2(BDSC-86108)

### Tumour generation in *Drosophila* imaginal discs

For generating epithelial tumours in larval imaginal discs, Mosaic Analysis with a Repressible Cell Marker (MARCM) technique was used(Wu and Luo, 2006). Tumour clones were generated by crossing the males of desired genotypes with virgin females from the MARCM stock (*y w hsFLP tubGAL4 UAS-GFP; FRT40A GAL80/CyO-GFP; +/+*) to create *lgl^4^ yki^OE^* clones expressing alone or co-expressing with the desired transgene/mutant allele. MARCM clones of *lgl^4^ yki^OE^*were generated essentially following Singh et. al.(Singh et al., 2022) Briefly, the clones were induced by exposing the early third instar larvae (72 h after egg laying (AEL)) to heat shock in a water bath at 37°C for 3 minutes. The following fly genotypes were used:

1. *y w hsFLP tubGAL4 UAS-GFP; tubGAL80 FRT40A/ lgl^4^yki^OE^ FRT40A; +/+*
2. *y w hsFLP tubGAL4 UAS-GFP; tubGAL80 FRT40A/ lgl^4^yki^OE^ FRT40A; hsp70 RNAi/+*
3. *y w hsFLP tubGAL4 UAS-GFP; tubGAL80 FRT40A/ lgl^4^yki^OE^ FRT40A; hsf RNAi/+*
4. *y w hsFLP tubGAL4 UAS-GFP; tubGAL80 FRT40A/ lgl^4^yki^OE^ FRT40A; sima RNAi/+*
5. *y w hsFLP tubGAL4 UAS-GFP; tubGAL80 FRT40A/ lgl^4^yki^OE^ FRT40A; dfoxo RNAi/+*
6. *y w hsFLP tubGAL4 UAS-GFP; tubGAL80 FRT40A/ lgl^4^yki^OE^ FRT40A; nox RNAi/+*
7. *y w hsFLP tubGAL4 UAS-GFP; tubGAL80 FRT40A/ lgl^4^yki^OE^ FRT40A; UAS-bsk^DN^/+*

### Treatment of the larvae with different inhibitors

500 μl yeast paste was prepared with 0.2g yeast powder, 10 μl food color (Green) and 200 mM of Pifithrin-μ (PFT-μ), Capivasertib [35 nM], N-acetyl cysteine (NAC) [2mg/ml] (Sigma-Aldrich, USA) or Butathione Sulfoximine (BSO) [200 mM]. Yeast paste was placed at the centre of Agar-petri plates. 20 *lgl^4^ yki^OE^* (48 h ACI) tumour bearing larvae were transferred to each petri-plates and kept at 24±1°C for 24 h or 48 h, followed by dissection of wing imaginal discs. In case of survivability analysis, larvae were fed with the drugs from 48 h ACI till they pupated and the pupae were followed till either their death as pupae or emergence as adult flies. Respective control flies were fed with yeast paste without PFT-μ, Capivasertib, NAC or BSO supplementation, respectively.

### Immunostaining and Confocal Microscopy

Imaginal discs of third instar larvae of appropriate genotypes and age (expressed as ACI for MARCM lines and days for other genotypes, as indicated in legends to figures) were dissected out in 1X Phosphate Buffer Saline (1X PBS) and immunostained following standard protocol(Ray et al., 2019). The primary antibodies used for experiments were mouse anti-Hsp83 (1:50,3E6, a kind gift from Prof. Robert Tanguay) (Morcillo et al. 1993); rat anti-Hsp70 (7Fb 1:200, Sigma), rabbit anti-Hsp60 (1:50, CST-SP60, D307#4870), rabbit anti-PH3, mouse anti-MMP1 (1:50, DSHB-14a3D2), rabbit anti-Dcp1 (1:100, Cell Signaling, Asp-216); rabbit anti-pJNK (1:400, V7932 Promega, USA). rat anti-Hsp/Hsc70 (1:50, 3.701, a kind gift from Prof. S. Lindquist) (Velazquez and Lindquist, 1984), rabbit anti-HSF (a kind gift from Prof. J. T. Lis; 1:200), rabbit anti-dFoxo (1:200, ab195977, Abcam). Appropriate secondary antibodies conjugated with Alexa Fluor 546 or Alexa Fluor 633 (1:200, Molecular Probes, USA) were used. To stain DNA and F-actin, 6-diamidino-2-phenylindole dihydrochloride (DAPI, 1 μg/ml) and Phalloidin-633 (Atto) were used, respectively. Immunostained tissue were mounted with 1,4-Diazabicyclo [2.2.2] octane (DABCO) anti-fade mounting medium. All the immunostained images were collected with Zeiss LSM 900 confocal microscope using Plan-10X (0.45 NA), Apo 20X (0.8 NA) or 63X oil immersion (1.4 NA) objectives and Zeiss ZEN 3.4 Blue software. Quantitative analysis of the images was carried out using Fiji (Image-J). All the images shown here were assembled using Adobe Photoshop (2012). Statistical analysis of control and experimental samples were done through unpaired t-test in GraphPad Prism 8.4.2.

### Survival assay

Larvae of the desired genotypes were allowed to develop till at 24±1°C. Numbers of those dying at early pupal (pseudopupae) or late pupal (pharate) stages and those that emerged as adults were counted. GraphPad Prism 8.4.2 was used to check the statistical significance between the different datasets using Student’s t-Test.

### Transplantation of tumour discs

GFP positive tumour clone bearing wing imaginal disc from *lgl^4^ yki^OE^* and *lgl^4^ yki^OE^*; *hsp70 RNAi* larvae (96 h ACI) or GFP-positive imaginal disc from Nub*-Gal4>UAS-GFP* larvae (3^rd^ instar) were dissected out in 70% Grace’s Insect medium. Discs were fragmented with the help of insulin needle using a GFPscope (Vaiseshika/Plan 1X) and one fragment/fly was introduced into the abdomen of 2 days old virgin *Oregon R+* females using a pulled glass capillary needle(Woodhouse et al., 1998). Injected host flies were kept in a food vial at 24±1°C for further observation till 14 days. Growth of GFP+ tumour and GFP+ control tissue within the host abdomen was imaged using a Nikon DS-Fi1c camera on a Nikon E800 Fluorescence microscope. 14 days post-transplantation, the host abdomens were dissected and immunostained with anti-Hsp70 and Phalloidin-633 as mentioned above.

### Imaging of Adult flies

Photographs of tumour patches in different body parts of surviving adults and abdomen morphology of transplanted host flies were taken with Axio Cam Zeiss camera mounted on Nikon SMZ800N stereo-binocular microscope.

### Analysis of RNAseq data

RNASeq data from an earlier published dataset (PMC id: PMC7768408) was reanalyzed to prepare a heat map showing differential expression of genes by SR plot (https://www.bioinformatics.com.cn/en). Gene sequence enrichment analysis was performed using Pangea.com. Significantly expressed genes were defined as those exhibiting an absolute fold-change of at least 1.5 and an FDR of less than or equal to 0.05.

## Acknowledgement

We heartily thank Dr. Bama Charan Mandal (Department of Zoology, BHU) and ISLS-BHU for Confocal microscope imaging system. We thank FIST-lab Zoology Dept. BHU for the central instrumentation facility. We heartily thank Prof. Pradip Sinha (IIT Kanpur, India) and Bloomington Drosophila Stock Centre (USA) for sharing the fly stocks with us. We share heartfelt gratitude to Prof. John. T. Lis (USA), Prof. Robert Tanguay (Canada), late Prof. S. Lindquist for sharing different antibodies, as mentioned in Materials and Methods.

## Author contributions

G.S., D.S., S.C.L. conceived the study. G.S. and A.B. designed experiments. Following generation of initial data by G.S., A.B. performed further experiments and analysed the data.

V.A. performed the meta-analysis of TCGA datasets. D.S. framed the manuscript and wrote it with the help of S.C.L and A.B.. D.S. and S.C.L. funded the research.

## Data availability

The data that support the findings of this study are available within the main text and its Supplementary Information file. The lead contact will share all raw data associated with this paper upon reasonable request and, when applicable, fulfilment of appropriate material transfer agreements.

