## Supplementary Fig S1 - S6 for "Non-canonically regulated heterochronic expression of Hsp70 drives clonal expansion and invasion in *Drosophila* epithelial tumours"

### Figure S1

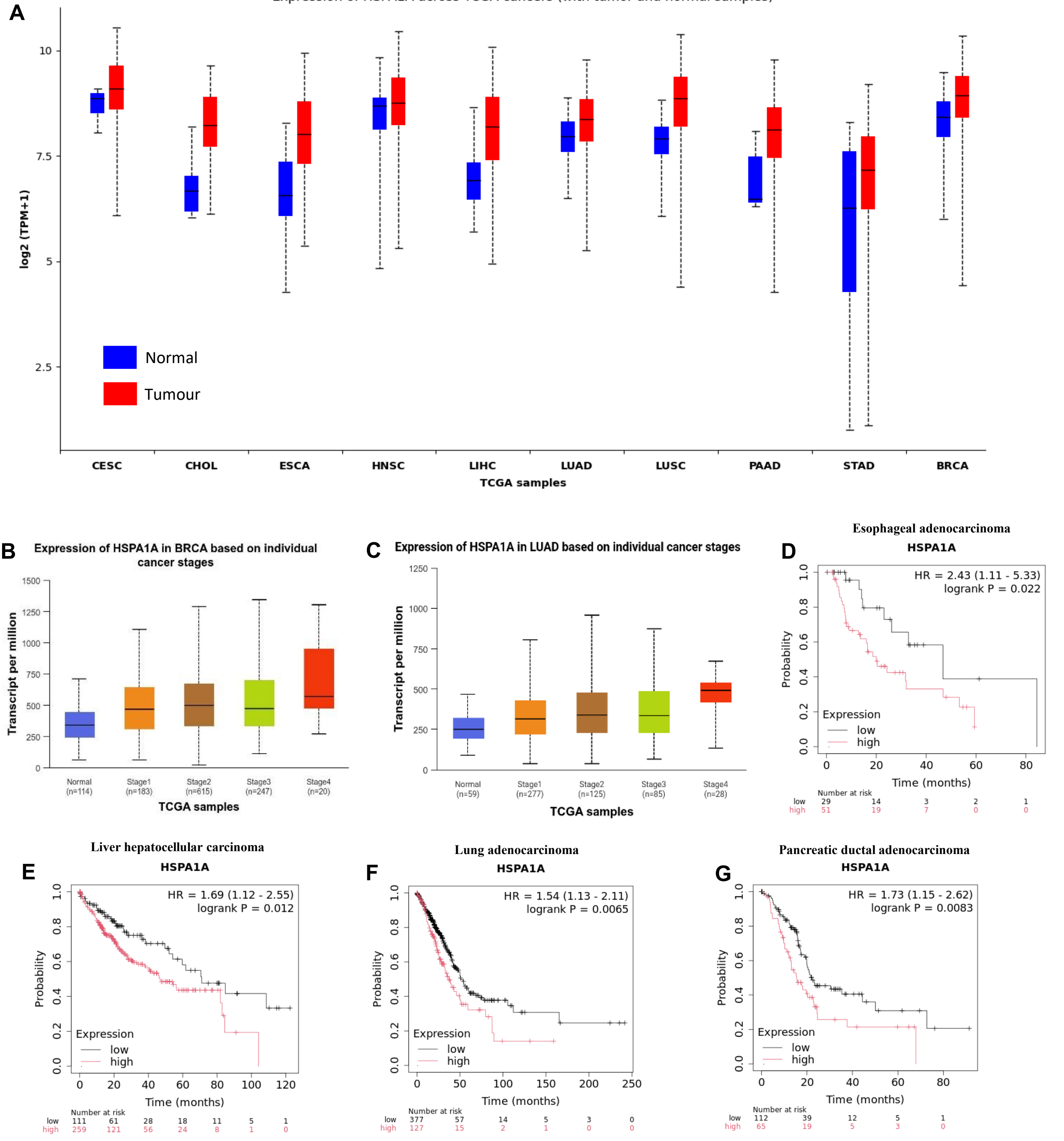
A
Expression of HSPA1A across TCGA cancers (with tumor and normal samples)
log2 (TPM+1)
Normal
Tumour
CESC
CHOL
ESCA
HNSC
LIHC
LUAD
LUSC
PAAD
STAD
BRCA
TCGA samples
B
Expression of HSPA1A in BRCA based on individual cancer stages
Transcript per million
Normal (n=114)
Stage1 (n=183)
Stage2 (n=615)
Stage3 (n=247)
Stage4 (n=20)
TCGA samples
C
Expression of HSPA1A in LUAD based on individual cancer stages
Transcript per million
Normal (n=59)
Stage1 (n=277)
Stage2 (n=125)
Stage3 (n=85)
Stage4 (n=28)
TCGA samples
D
Esophageal adenocarcinoma
HSPA1A
HR = 2.43 (1.11 - 5.33)
logrank P = 0.022
Probability
Expression
low
high
Time (months)
Number at risk
low 29 14 3 2 1
high 51 19 7 0 0
E
Liver hepatocellular carcinoma
HSPA1A
HR = 1.69 (1.12 - 2.55)
logrank P = 0.012
Probability
Expression
low
high
Time (months)
Number at risk
low 111 61 28 18 11 5 1
high 259 121 56 24 8 1 0
F
Lung adenocarcinoma
HSPA1A
HR = 1.54 (1.13 - 2.11)
logrank P = 0.0065
Probability
Expression
low
high
Time (months)
Number at risk
low 377 57 14 5 3 0
high 127 15 2 1 0 0
G
Pancreatic ductal adenocarcinoma
HSPA1A
HR = 1.73 (1.15 - 2.62)
logrank P = 0.0083
Probability
Expression
low
high
Time (months)
Number at risk
low 112 39 12 5 1
high 65 19 5 3 0

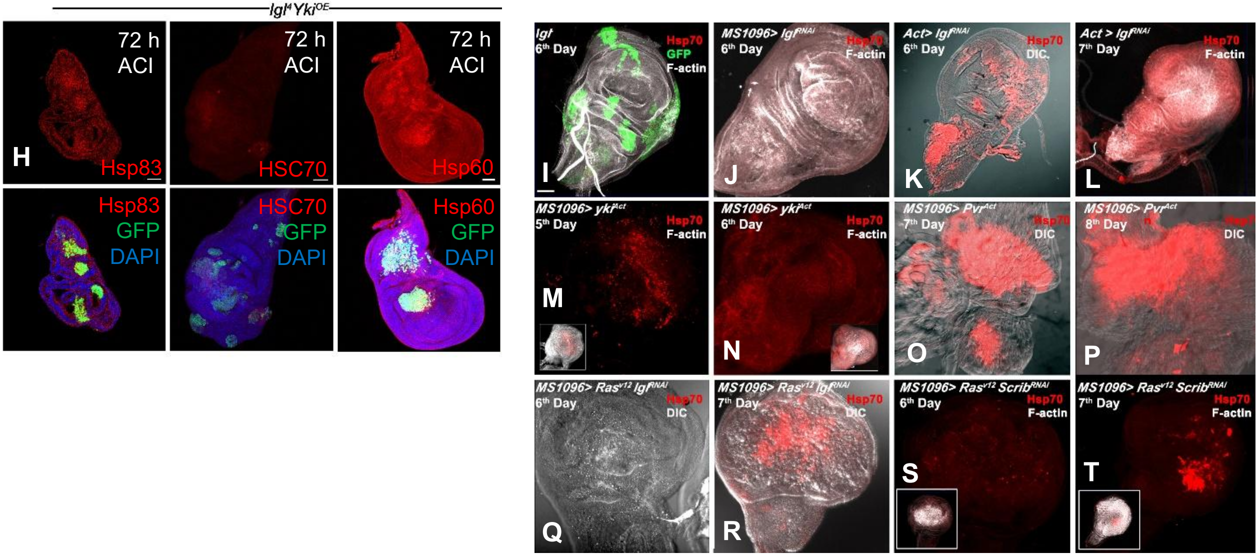


**Fig S1.** Analysis of HspA1A (Hsp70) levels across The Cancer Genome Atlas database and different *Drosophila* tumour genotypes. (**A**) Shows differential expression analysis of HspA1A between tumor and normal samples across various cancer types using the TCGA database, Cancer type abbreviations: Cervical Squamous cell carcinoma (CESC), Cholangiocarcinoma (CHOL), Esophageal carcinoma (ESCA), Head and neck squamous cell carcinoma (HNSC), Liver hepatocellular carcinoma (LIHC), Lung adenocarcinoma (LUAD), Lung squamous cell carcinoma (LUSC), Pancreatic adenocarcinoma (PAAD), Stomach adenocarcinoma (STAD), Breast invasive carcinoma (BRCA). (**B-C**) Analyses the stage-specific expression of HspA1A in (B) Breast invasive carcinoma (BRCA) and (C) Lung adenocarcinoma (LUAD), using TCGA dataset. (**D-G**) Kaplan-Meier survival analysis demonstrating the effect of HspA1A expression on overall survival in different cancer types (D-Esophageal adenocarcinoma, E-Liver hepatocellular carcinoma, F-Lung adenocarcinoma, G-Pancreatic ductal adenocarcinoma) from the Kaplan-Meier Plotter. (**H**) Panel showing expression of other Heat Shock Proteins in $lgl^4$ $yki^{OE}$ tumor background, Scale bar: 50µm. (**I-T**) Confocal projection images of wing discs (genotypes noted on the top left corner in each case) showing Hsp70 (red) and F-actin (white) staining, with or without DIC (gray) image; panel in (I) also shows GFP (green) in the $lgl^-$ MARCM clones at 96 h ACI; larval age is indicated on upper left corner of each panel as days after egg laying. Insets in (M, N, S and T) show low magnification images indicating F-actin and Hsp70 staining, of the respective wing discs showing only Hsp70 (red) staining. Scale bar in (I) represents 50µm and applies to I-T.

**Figure S2**

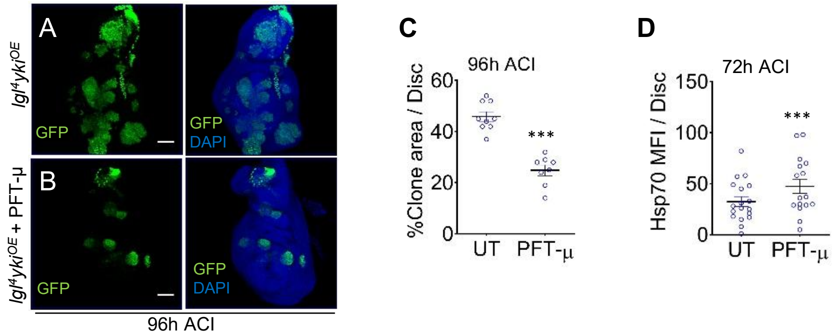


**Fig S2.** Effect of PFT-μ mediated inhibition of Hsp70 in *lgl*$^4$ *yki*$^{OE}$ tumour size. (**A-B**) Confocal projection images showing GFP+ clones in untreated (UT) *lgl*$^4$ *yki*$^{OE}$ and PFT - μ treated *lgl*$^4$ *yki*$^{OE}$ tumour wing disc (96 h ACI and 48 h after drug treatment (ADT)), Scale bar 50 μm. (**C**) Graph showing the clonal area of PFT - μ treated larval wing discs bearing *lgl*$^4$ *yki*$^{OE}$ clone compared to untreated ones (n =9), ***$p < 0.001$, unpaired Student's t- test. (**D**) Plot showing the mean intensity of Hsp70 labelling per clone in PFT-μ treated *lgl*$^4$ *Yki*$^{OE}$ tumour discs compared to untreated, at 72 h ACI or 24 h ADT) (n = 20), ***$p < 0.001$, unpaired Student's t- test.

**Figure S3**

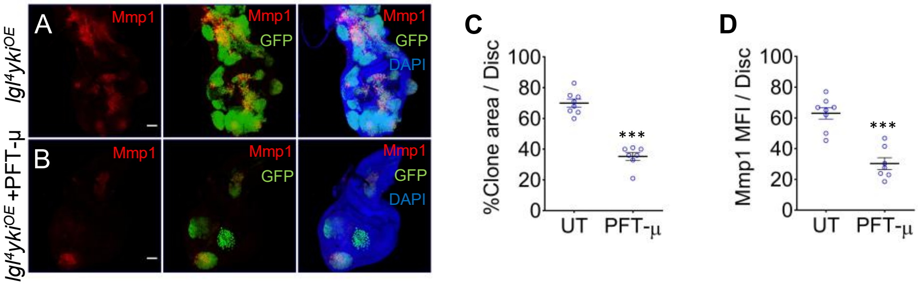


**Fig S3.** Effect of PFT-μ mediated inhibition of Hsp70 in *lgl*$^{4}$ *yki*$^{OE}$ tumour on MMP1 expression. (**A-B**) Confocal projection images of wing disc bearing *lgl*$^{4}$ *yki*$^{OE}$ (untreated (UT)) and *lgl*$^{4}$ *yki*$^{OE}$ (PFT-μ treated) clones showing MMP1 in clonal area (green) at 96 hr ACI + 48h ADT, Scale bar 50 μm. (**C**) Quantification plot representing clonal area of PFT-μ treated larval wing discs bearing *lgl*$^{4}$ *yki*$^{OE}$ clone, compared to untreated (UT) *lgl*$^{4}$ *yki*$^{OE}$ clone bearing discs at 48h after treatment (96h ACI)**,** (n = 8), ***p < 0.001, unpaired Student's t- test. (**D**) Mean MMP1 signal intensity per clone in *lgl*$^{4}$ *yki*$^{OE}$ (PFT- μ treated) clone bearing discs compared to *lgl*$^{4}$ *yki*$^{OE}$ control clones, (n = 8) ***p < 0.001, unpaired Student's t- test.

**Figure S4**

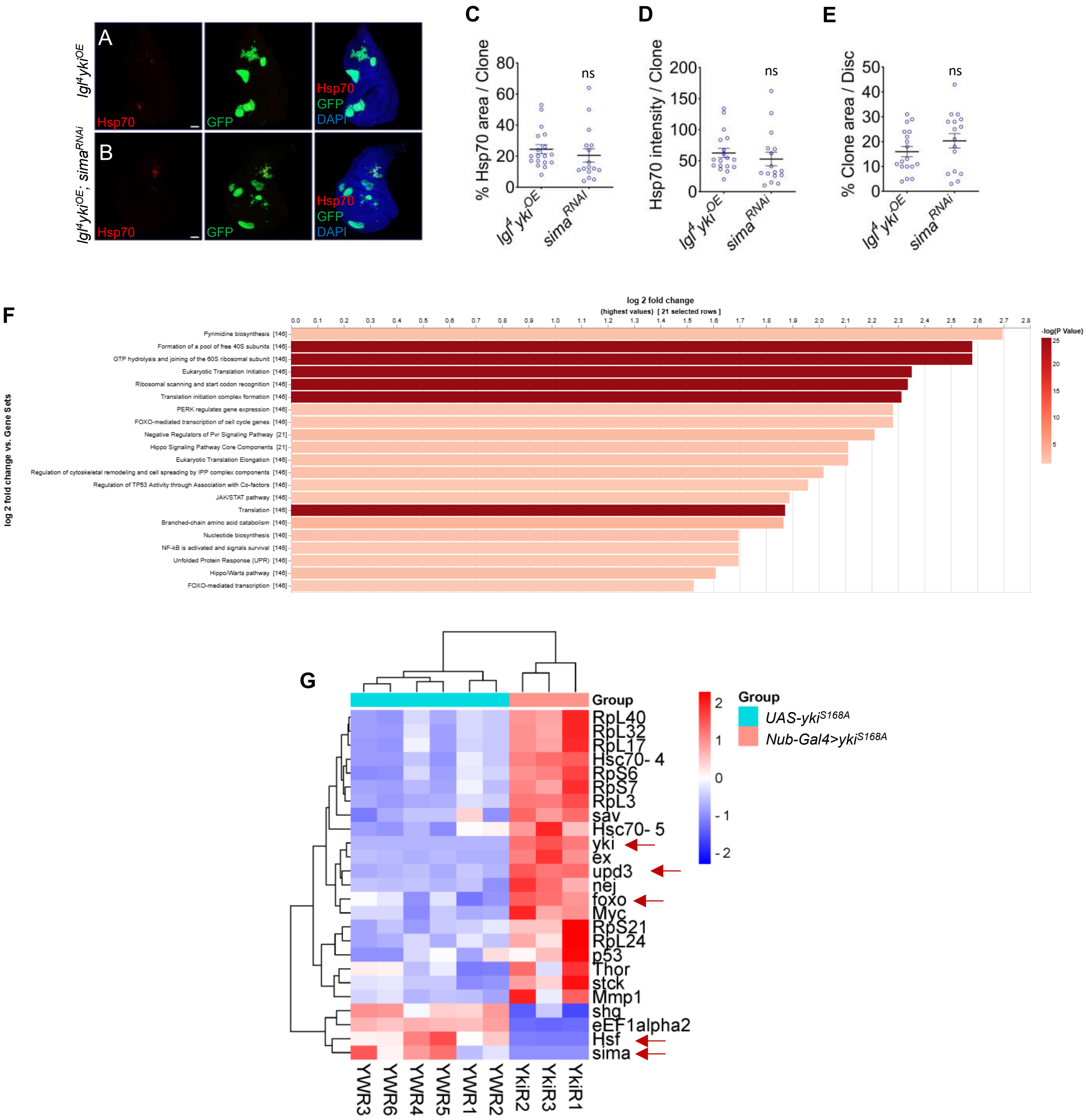
A
B
lgl4 ykiOE
lgl4 ykiOE; simaRNAi
Hsp70
GFP
DAPI
C
D
E
% Hsp70 area / Clone
Hsp70 intensity / Clone
% Clone area / Disc
ns
lgl4 ykiOE
simaRNAi
F
log 2 fold change
(highest values) [ 21 selected rows ]
-log(P Value)
log 2 fold change vs. Gene Sets
Pyrimidine biosynthesis [146]
Formation of a pool of free 40S subunits [146]
GTP hydrolysis and joining of the 60S ribosomal subunit [146]
Eukaryotic Translation Initiation [146]
Ribosomal scanning and start codon recognition [146]
Translation initiation complex formation [146]
PERK regulates gene expression [146]
FOXO-mediated transcription of cell cycle genes [146]
Negative Regulators of Pvr Signaling Pathway [21]
Hippo Signaling Pathway Core Components [21]
Eukaryotic Translation Elongation [146]
Regulation of cytoskeletal remodeling and cell spreading by IPP complex components [146]
Regulation of TP53 Activity through Association with Co-factors [146]
JAK/STAT pathway [146]
Translation [146]
Branched-chain amino acid catabolism [146]
Nucleotide biosynthesis [146]
NF-kB is activated and signals survival [146]
Unfolded Protein Response (UPR) [146]
Hippo/Warts pathway [146]
FOXO-mediated transcription [146]
G
Group
UAS-ykiS168A
Nub-Gal4>ykiS168A
RpL40
RpL32
RpL17
Hsc70- 4
RpS6
RpS7
RpL3
sav
Hsc70- 5
yki
ex
upd3
nej
foxo
Myc
RpS21
RpL24
p53
Thor
stck
Mmp1
shg
eEF1alpha2
Hsf
sima
YWR3
YWR6
YWR4
YWR5
YWR1
YWR2
YkiR2
YkiR3
YkiR1

**H**

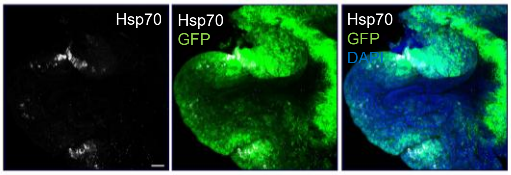


*Nubbin-Gal4>UAS-ykiS168A*

**Fig S4.** Hsp70 expression is independent of Sima. (**A-B**) Confocal projection images showing GFP+ clonal area and Hsp70 expression in *lgl*$^{4}$ *yki*$^{OE}$ *and lgl*$^{4}$ *yki*$^{OE}$*; sima*$^{RNAi}$ clone bearing disc at 72 h ACI*,* Scale bar 50 µm. (**C-D**) Quantification plot showing insignificant difference in Hsp70 labeled area (C) and Hsp70 intensity per clone (D) between *lgl*$^{4}$ *yki*$^{OE}$ and *lgl*$^{4}$ *yki*$^{OE}$ *; sima*$^{RNAi}$ clone bearing discs (n = 17), ns – non-significant, $p > 0.05$ , unpaired Student's t- test. (**E**) Quantification plot showing insignificant differences in clonal area per disc between *lgl*$^{4}$ *yki*$^{OE}$ and *lgl*$^{4}$ *yki*$^{OE}$*; sima*$^{RNAi}$ clone bearing discs at 72hr ACI (n = 17), ns – non-significant, $p > 0.05$, unpaired Student's t-test. (**F**) Heat map representing the expression level of differentially expressed genes between wild type (*UAS-Yki*$^{S168A}$) and *Nub-Gal4>Yki*$^{S168A}$ group, filtered based on flog2 fold >1.5, $p_{adj}<0.05$, and false discovery rate (FDR) lower than 15%. (**G**) Bar chart representing enriched pathways ($p < 0.05$) identified using Pangea for *Yki*$^{S168A}$ overexpressing discs compared to control. (**H**) Confocal projection images depicting Hsp70 expression pattern (white) in *Nubbin-Gal4>UAS-ykiS168A* tumours (green).

**Figure S5**

**A**

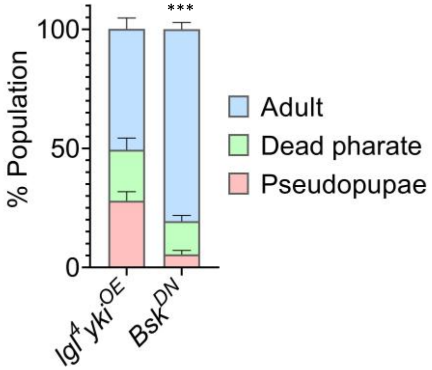


**Fig S5.** Survival of *lgl⁴ ykiOE; bskDN* flies. (**A**) Stacked bar graph comparing mean % (+ S.E.) of pseudopupae, pharate and adult survivors of *lgl⁴ ykiOE* and *lgl⁴ ykiOE; bskDN* tumour backgrounds, (n = 237 larvae of each genotype) $*p < 0.05$, $***p < 0.001$, unpaired Student's t- test.

**Figure S6**

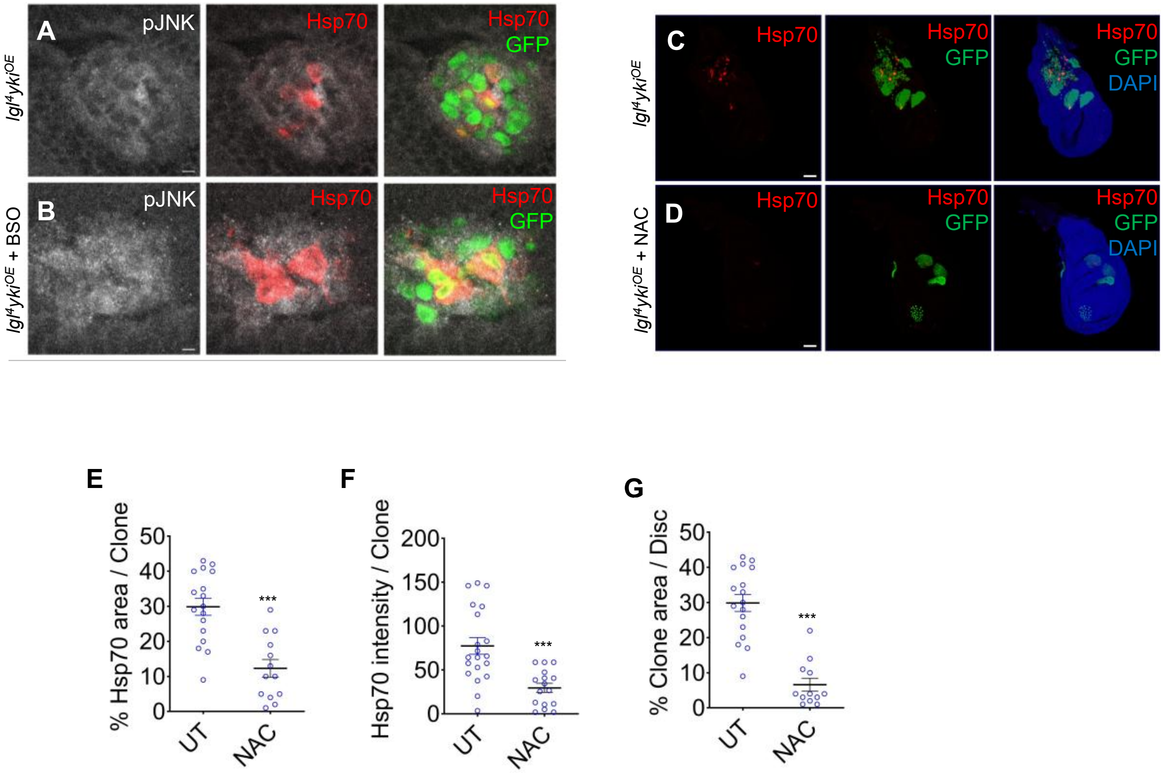


**Fig S6.** Alteration of ROS affects Hsp70 expression in *$lgl^4$ $Yki^{OE}$* clones**.** (**A-B**) Magnified confocal optical section showing more colocalization of pJNK and Hsp70 in BSO treated *$lgl^4$ $yki^{OE}$* clone bearing disc than untreated *$lgl^4$ $yki^{OE}$* disc, Scale bar 50µm. The corresponding quantitative analysis shown in Fig 6M. (**C-D**) Confocal projection image showing expanded GFP+ clonal area and high Hsp70 expression in *$lgl^4$ $yki^{OE}$* untreated (UT) larval disc than that of NAC treated ones, Scale bar 50 µm. (**E**) Quantification plot showing % of clonal area (green) covered by Hsp70, in NAC treated *$lgl^4$ $yki^{OE}$* larval clonal disc (n = 13) verses untreated controls (n = 17) ***p < 0.001, unpaired Student's t- test. (**F**) Quantification plot showing Hsp70 signal intensity in clonal area (green) of *$lgl^4$ $yki^{OE}$* tumour bearing NAC treated discs (n = 13) verses untreated controls (n = 17), ***p < 0.001, unpaired Student's t- test. (**G**) Plot showing % of clonal area (green) of *$lgl^4$ $yki^{OE}$* (NAC treated) clone bearing discs (n = 13) verses *$lgl^4$ $yki^{OE}$* bearing discs (n = 17) ***p < 0.001, unpaired Student's t- test.
